# Cycling Molecular Assemblies (CyMA) for Ultrasensitive Golgi Imaging

**DOI:** 10.1101/2025.11.19.689330

**Authors:** Weiyi Tan, Qiuxin Zhang, Erica C. Dresselhaus, Divyanshu Mahajan, Isabela Ashton-Rickardt, Lei Lu, Avital A. Rodal, Bing Xu

## Abstract

The Golgi apparatus is central to intracellular trafficking, yet its dynamic visualization remains constrained by probes that require high concentrations and long incubations. Here we present cycling molecular assemblies (CyMA), a design concept that harnesses endogenous dynamic enzymatic futile cycle to drive ultrasensitive imaging. The BODIPY-CyMA probe operates through reversible palmitoylation-depalmitoylation mediated by palmitoyl acyltransferases and thioesterases, establishing a nonequilibrium steady-state that actively concentrates the probe at the Golgi. This enzymatic cycling converts diffusion-limited localization into self-amplifying signal generation, enabling morphology imaging at concentrations as low as 100 pM and real-time tracking of Golgi dynamics within minutes at nanomolar levels. This probe requires minimal incubation time, exhibits negligible cytotoxicity, faithfully reports pharmacological Golgi disassembly, and functions *in vivo* in *Drosophila* larvae. BODIPY-CyMA exemplifies how coupling molecular self-assembly to endogenous enzymatic cycles affords a general strategy for constructing dynamic, non-perturbative probes for live cell and *in vivo* imaging.

## Introduction

This article reports on the use of cycling molecular assemblies (CyMA) for ultrasensitive Golgi imaging. Golgi apparatus serves as a central hub for post-translational modification, sorting, trafficking of proteins and lipids, and its dynamic morphology reflects fundamental aspects of cellular regulation.^[1]^ Real-time visualization of these morphological rearrangements, particularly the interactions of Golgi with other cellular components, provides deeper insights into fundamental cellular mechanisms beyond static structural snapshots or proteogenomic data.^[2]^ Such dynamic imaging would elucidate the role of Golgi in maintaining cellular homeostasis and modulating signaling pathways, offering a more comprehensive knowledge on the context-dependent roles of Golgi in cell biology. Consequently, there has been steady effort^[3]^ to develop more sensitive and versatile imaging probes for understanding Golgi functions since the introduction of the classical trans-Golgi probe C6-NBD-ceramide.^[4]^ Beyond small molecules, fluorescent proteins are widely used for live-cell imaging of the Golgi due to their ability to provide real-time visualization of cellular processes, despite potential limitations such as photobleaching, functional interference, and the need for genetic transfection.^[5]^

Despite progress in Golgi imaging probes, significant challenges remain. Small molecule probes usually need high micromolar concentrations or prolonged incubation time^[6]^, which is often incompatible with minimally perturbative imaging or the study of rapid cellular processes. Beyond improving performance metrics like brightness, a key unmet need is for probes that report on the physiological state of the Golgi itself. The identity and function of the Golgi are actively maintained by a unique landscape of resident enzymes. An ideal probe, therefore, would not merely stain the organelle but would instead engage with this specific enzymatic machinery, making its localization and signal a direct reflection of the dynamic functional activity of Golgi. However, achieving this requires probes that operate at ultralow concentrations to avoid the potential artifacts and perturbations associated with high doses of lipophilic molecules.

Our recent study uncovered that NBD-containing thiophosphopeptides^[7]^ and peptide thioesters^[8]^ efficiently target the Golgi, with the probe concentrations of 500 nM and 2 µM, respectively. Notably, these probes can localize at the Golgi almost instantly, requiring minimal incubation time. Our mechanistic studies revealed that these molecules undergo enzymatic reactions to form thiopeptides. The thiopeptides, undergoing reversible deacylation and acylation at the Golgi via thioesterase (TE) and palmitoyl acyltransferase (zDHHC) catalysis, generate CyMA, non-diffusible, yet dynamic molecular assemblies,^[9]^ for Golgi imaging. With the first-in-class molecular mechanism, CyMA provide a framework for fast and sensitive Golgi imaging.

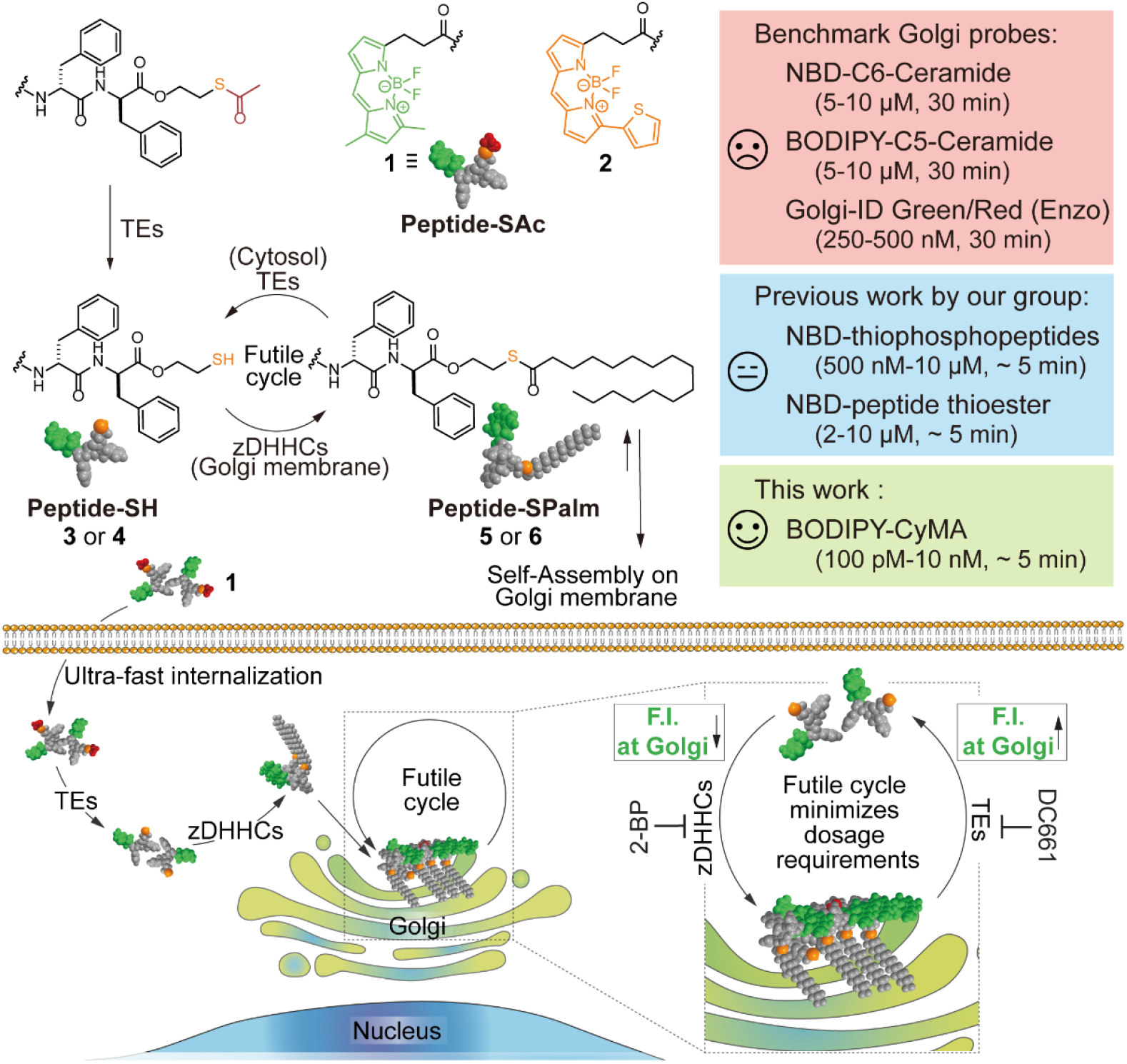

Building on mechanistic insights of CyMA^[9]^ and the superior photophysical properties of BODIPY,^[3a, b]^ we conjugated BODIPY to peptide thioesters to generate BODIPY-CyMA precursors (**1** and **2**) with distinct excitation wavelength (**Scheme 1**). These BODIPY-incorporated CyMA accumulate at the Golgi even at picomolar concentrations via reversible S-acylation catalyzed by TEs and zDHHCs. The resulting enzymatic futile cycle continuously recycles diffused intermediates, enriching the local probe concentration and enabling effective Golgi labeling at ultralow doses. Moreover, 50 nM of **2** is sufficient to image the Golgi in *drosophila* larvae within 10 minutes. Furthermore, the co-assembly of **2** with its FRET quencher UBQ2-CyMA (**7**) results in fluorescence intensity reduction at the Golgi, supporting the formation of CyMA within this organelle. Importantly, both **1** and **2** exhibit excellent biocompatibility, with no detectable cytotoxicity (**Figure 3D**). These properties establish CyMA as a highly effective probe for imaging Golgi dynamics in live cells, as demonstrated by real-time observations of structural and functional disruptions induced by brefeldin A (BFA)^[2a]^ and nocodazole^[10]^. By leveraging the enzymatic S-lipidation cycles regulated by deacylases and acylases, CyMA achieve effective Golgi targeting at sub-nanomolar concentrations, representing a substantial advance over the aforementioned conventional Golgi probes. This imaging strategy not only overcomes the limitations of existing Golgi probes but also opens new avenues for studying the dynamic functions of the Golgi with both high sensitivity and fast speed.

## Results and Discussion

### BODIPY-CyMA precursor design and synthesis

The core design of BODIPY-CyMA precursors integrates the BODIPY fluorophore with essential structural elements that enable dynamic, enzyme-mediated accumulation within the Golgi^[7a, 8-9]^. These structural features include: (i) a self-assembling peptide motif (i.e., D-Phe-D-Phe^[11]^), which promotes supramolecular organization, as confirmed by recent cryo-EM structures^[12]^; (ii) a strategically placed thioester warhead, which responds to intracellular enzymatic activities involving thioesterases (TEs) and palmitoyl acyltransferases (zDHHCs), thereby engaging in futile cycles of reversible S-palmitoylation (**Scheme 1**) that drive selective accumulation and dynamic assembly at the Golgi; and (iii) a well-positioned ester bond in the linker region, which represents a rational design strategy to improve membrane permeability in peptide-based constructs^[13]^, thereby enhancing cellular uptake. We decided to use D-Phe-D-Phe because aromatic residues were essential for Golgi targeting.^[8]^

The differences in functional group reactivity allows efficient synthesis of BODIPY-CyMA (Scheme S1). Initially, mercaptoethanol selectively attaches to 2-chlorotrityl resin through its thiol group under base-free dichloromethane (DCM) conditions. Steglich esterification connects the exposed hydroxyl group to Fmoc-D-phenylalanine, followed by standard Fmoc-based solid-phase peptide synthesis (SPPS) to assemble the desired peptide sequence. Cleavage from the resin using trifluoroacetic acid (TFA) simultaneously exposes the thiol, which reacts *in situ* with acyl chloride to form a thioester linkage. Under these acidic conditions, the N-terminal amine is protonated, thereby suppressing its reactivity and ensuring selective thioester formation. Finally, the free N-terminal amine of the peptide thioester is coupled to an activated BODIPY-carboxylic acid, yielding the final BODIPY-CyMA precursors.

### BODIPY-CyMA enables Golgi labeling at ultralow concentrations

As shown in **Figure 1A** and **1C**, BODIPY-CyMA precursor (**1** or **2**) retains its Golgi-targeting capability even at ultralow concentrations. Following a 4-h treatment with **1** or **2** at 10 nM, we observed bright Golgi fluorescence. Remarkably, as the concentrations decreased from 10 nM to 100 pM, the Golgi morphology remained clearly visible even at 100 pM. Quantification of the Golgi fluorescence intensity (**Figure 1B** and **1D**) further supports the excellent Golgi labeling performance of BODIPY-CyMA at these ultralow concentrations. For comparison, 10 nM BODIPY-C5-Ceramide exhibited very dim fluorescence at the Golgi, comparable to that observed in cells treated with 500 pM of **1** (**Figure S1**), indicating over a 20-fold enhancement in sensitivity for Golgi labeling with BODIPY-CyMA. Moreover, BODIPY-C5-Ceramide showed no detectable Golgi localization at picomolar concentrations (**Figure S1A**), further highlighting the superior performance of BODIPY-CyMA as a Golgi probe.

**Figure 1.**
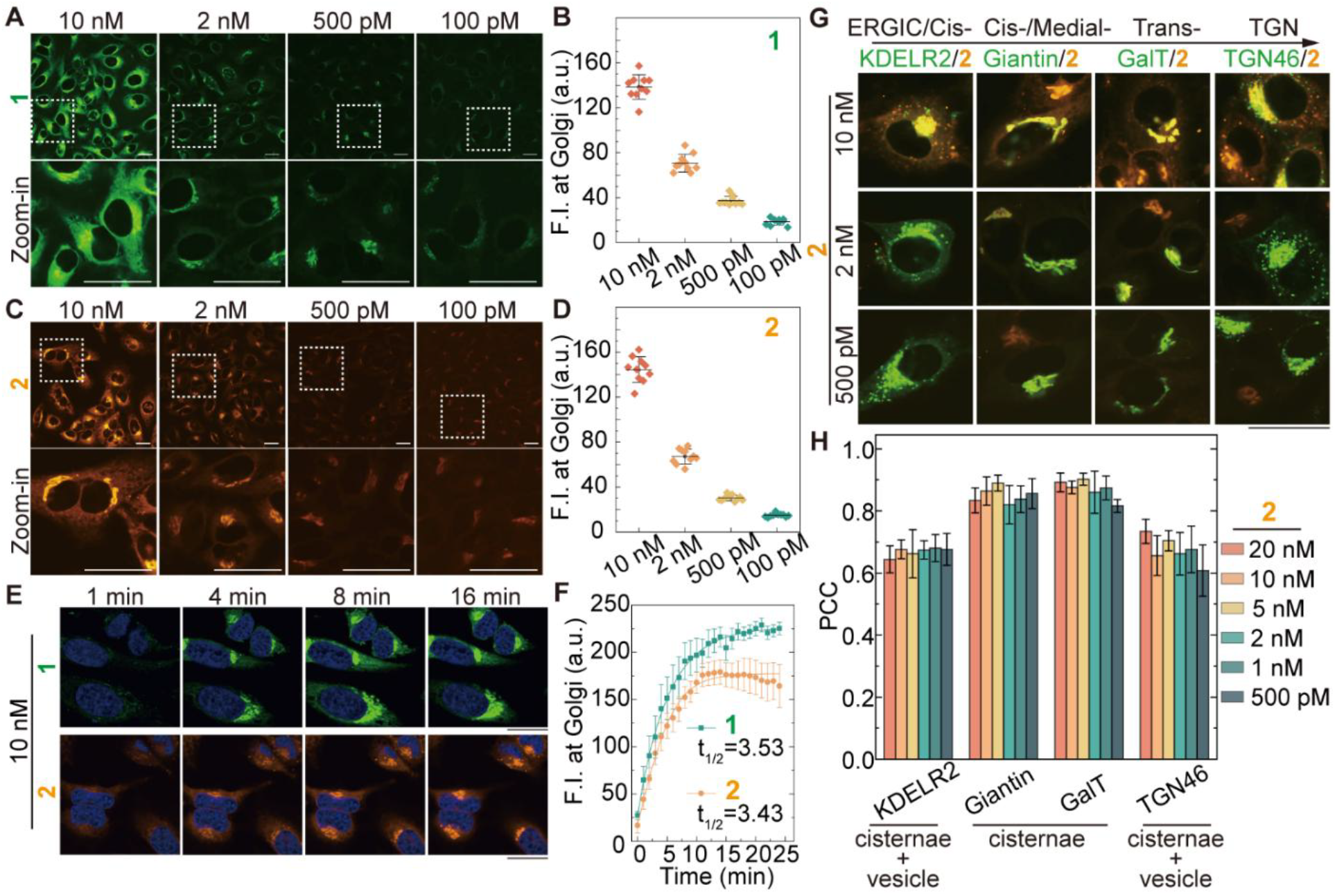
Ultrafast Golgi localization of BODIPY-CyMA at ultra-low concentrations. (A) CLSM of HeLa cells treated with various concentrations of **1** for 4 hours. (B) Quantitative analysis of fluorescence intensity at Golgi of HeLa cells in (A) (n = 10). (C) CLSM of HeLa cells treated with various concentrations of **2** for 4 hours. (D) Quantitative analysis of fluorescence intensity at Golgi of HeLa cells in (C) (n = 10). (E) Time-lapse CLSM images of HeLa cells treated with 10 nM of **1** or **2** over 24 minutes. (F) Quantitative analysis of fluorescence intensity build-up in HeLa cells treated with 10 nM of **1** or **2** over 24 minutes, as well as t_1/2_ (n = 10). (G) Colocalization study of various concentrations of **2** with Golgi markers (KDELR2, ERGIC/cis-Golgi; Giantin, cis-/medial-Golgi; GalT, trans-Golgi; TGN46, trans-Golgi network). (H) Pearson’s correlation coefficient (PCC) of **2** with Golgi markers (n = 5).

### BODIPY-CyMA exhibits ultrafast Golgi labeling

To assess the labeling kinetics, we evaluated the Golgi accumulation of BODIPY-CyMA at a concentration of 10 nM. Fluorescence was built up quickly at the perinuclear region within 1 minute, and we can clearly observe the Golgi morphology and localization with 2 minutes (**Figure 1E, Movie S1**). Both BODIPY-CyMA precursors **1** and **2** demonstrated an ultrafast Golgi accumulation, with half-times (t_1/2_) of approximately 3.5 minutes (**Figure 1F**). In contrast, no significant Golgi fluorescence was detected in cells treated with 10 nM BODIPY-C5-Ceramide (**Figure S2**). Given that no known endocytic pathway is capable of transporting materials from the extracellular space to the Golgi within a t_1/2_ shorter than 5 minutes, it is plausible that the amphiphilic BODIPY-CyMA precursors bypass conventional endosomal routes and instead traverse the plasma membrane for rapid accumulation at the Golgi^[9]^.

### Specific Golgi labeling and preferential cisternal accumulation of BODIPY-CyMA

To further evaluate Golgi labeling specificity, we performed colocalization studies using GFP-tagged Golgi markers, including KDELR2 (ERGIC/cis-Golgi), Giantin (cis-/medial-Golgi),^[14]^ GalT (trans-Golgi), and TGN46 (trans-Golgi network, TGN) (**Figure 1G, Figure S3-S6**). We tagged the proteins with GFP because there is no crosstalk between GFP and **2**. Across a range of BODIPY-CyMA concentrations (20 nM to 500 pM), Pearson’s correlation coefficients (PCCs) with each Golgi marker remained consistent, indicating the stability and specificity of **2** in Golgi labeling. Notably, BODIPY-CyMA exhibited PCC values above 0.7 with KDELR2 and TGN46 and even higher values (above 0.85) with Giantin and GalT (**Figure 1H**). These results demonstrate not only the excellent Golgi-targeting capability of BODIPY-CyMA but also suggest a preferential localization within the Golgi cisternae. This interpretation is further supported by high-resolution CLSM imaging (**Figure S7**), which revealed KDELR2- and TGN46-positive vesicles, reflecting their known localization at the ERGIC-Golgi interface^[15]^ and the TGN exit^[16]^, respectively. BODIPY-CyMA hardly colocalize with these vesicular structures, suggesting that its fluorescence is largely restricted to the cisternal compartments. Given that Giantin and GalT are specifically localized within the Golgi cisternae^[17]^, the strong colocalization with these markers reinforces the conclusion that BODIPY-CyMA preferentially accumulates in the cisternal regions of the Golgi.

### BODIPY-CyMA precursor requires deacetylation and palmitoylation for Golgi targeting

To confirm that BODIPY-CyMA probes function through counteracting TEs and zDHHCs, we treated cells with TE inhibitors (ML211 for LYPLA1/2^[18]^ or DC661 for PPT1^[19]^) and observed that both inhibitors effectively reduced the Golgi accumulation of **1** and **2** (**Figure 2A, Figure S8**). This result suggests that the deacetylation of **1** or **2** to generate the corresponding peptide-SH species (**3** or **4**) is essential for Golgi targeting (**Figure 2B**). Subsequently, inhibiting palmitoylation by treating cells with the zDHHC inhibitor 2-bromopalmitate (2-BP) also caused a significant decrease in Golgi fluorescence (**Figure 2A, Figure S9**). Direct evidence for this modification in cells was obtained through liquid chromatography-high-resolution mass spectrometry (LC-HRMS), which confirmed the presence of the palmitoylated species in treated cells (**Figure S10**). These results demonstrate that the S-palmitoylation of the peptide-SH intermediate to form peptide-SPalm is crucial for the probe’s retention within the Golgi. Mechanistically, the addition of the C16 palmitoyl chain markedly increases the hydrophobicity of the probe, facilitating its self-assembly into the Golgi membrane, where zDHHC enzymes are embedded^[20]^.

**Figure 2.**
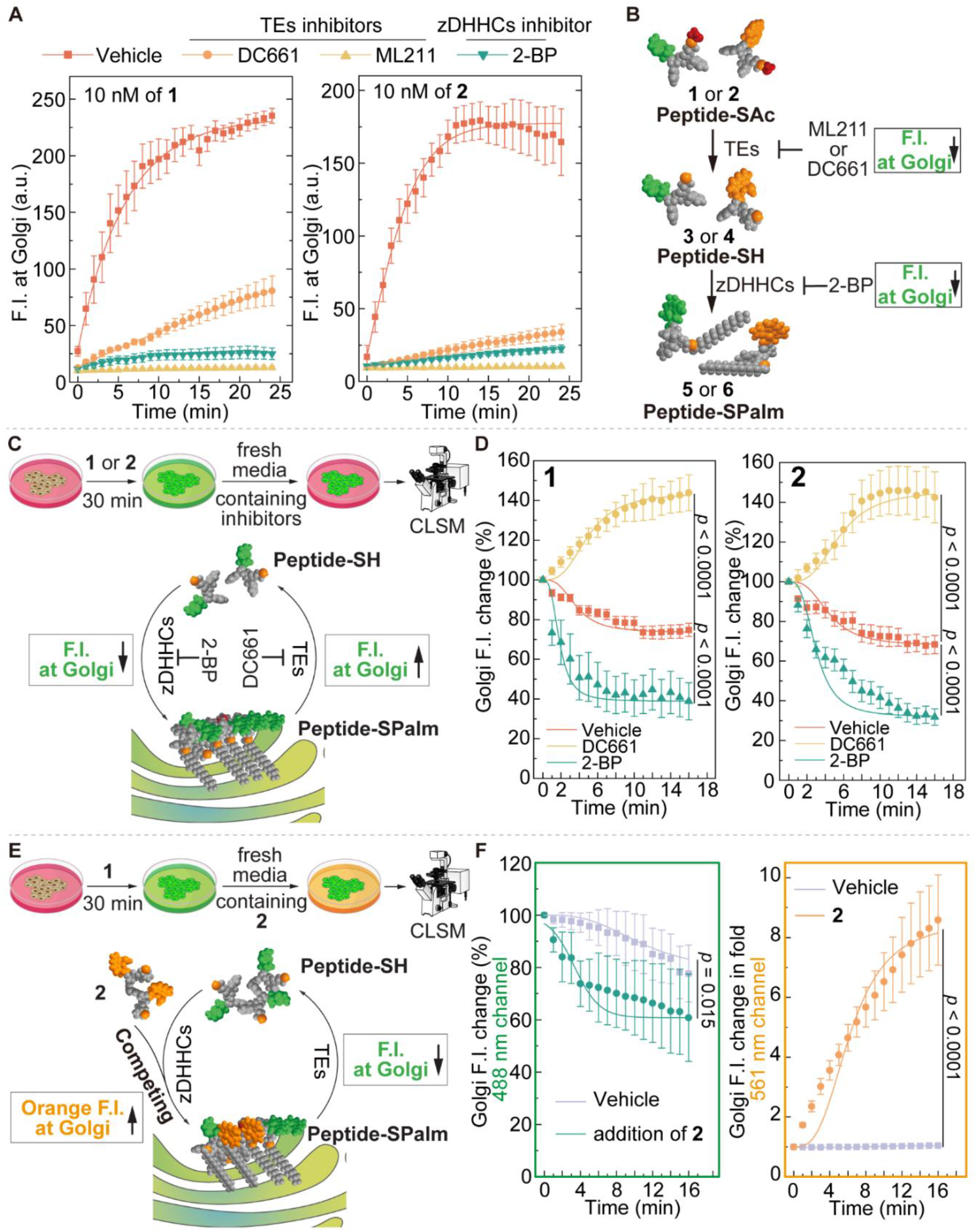
BODIPY-CyMA undergoes deacetylation and palmitoylation, generating futile cycles by palmitoylation/depalmitoylation reactions. (A) Golgi fluorescence intensity of HeLa cells pretreated with thioesterases inhibitor (DC661, 20 μM or ML211, 50 μM) or zDHHCs inhibitor (2-BP, 10 μM) for 30 minutes, and then incubated with **1** or **2** (10 nM) (n = 10). (B) Schematic illustration of studying the role of TEs and zDHHCs in Golgi targeting by BODIPY-CyMA precursors. (C) Schematic illustration of experimental design, cells were pretreated with **1** or **2** (10 nM, 30 min), then switched to fresh media with either PPT1 inhibitor (DC661, 20 μM) or zDHHC inhibitor (2-BP, 20 μM) and monitored the Golgi fluorescence intensity change. (D) Quantitative analysis of the Golgi fluorescence intensity changes of HeLa cells in (C) (n = 10). (E) Schematic illustration of experimental design, cells were pretreated with **1** (10 nM, 30 min), then switched to fresh media with **2** (10 nM, 30 min) and monitored the Golgi fluorescence intensity change in both 488 nm and 561 nm channels. (F) Quantitative analysis of the Golgi fluorescence intensity changes of HeLa cells in (E) in both channels (n = 10).

### BODIPY-CyMA establishes palmitoylation-depalmitoylation cycle at the Golgi

We further used two complementary approaches to validate the presence of a palmitoylation-depalmitoylation cycle at the Golgi (**Figure 2C,E**). In the first approach (**Figure 2C**), we pretreated cells with **1** or **2** for 30 minutes to establish a steady-state among the peptide-SAc, peptide-SH, and peptide-SPalm species. After washing out the precursors and replacing the medium with fresh media containing either TE inhibitor (DC661) or zDHHC inhibitor (2-BP), we monitored changes in Golgi fluorescence intensity. Upon removal of **1** or **2** in the culture medium, we observed a gradual decrease in fluorescence intensity at the Golgi, reflecting the disruption of the steady-state balance (**Figure 2D**). Incubating cells with 2-BP significantly accelerated this fluorescence decay (**Figure 2D, Figure S11**). This occurs because blocking *de novo* palmitoylation prevents the replenishment of the Golgi-localized peptide-SPalm pool from peptide-SH, while depalmitoylation continues. Conversely, treatment with DC661 increased Golgi fluorescence (**Figure 2D, Figure S11**), indicating that blocking TEs shifts the balance toward peptide-SPalm, thereby enhancing Golgi accumulation. Together, the opposing effects of inhibiting palmitoylation versus depalmitoylation provide strong evidence for the proposed enzymatic cycle governing BODIPY-CyMA retention at the Golgi.

The second approach is chasing **1** with **2** (**Figure 2E**), because of their distinctive excitation/emission spectrum but sharing the same Golgi targeting mechanism. To be more specific, we first incubated cells with **1** to establish steady-state green fluorescence at the Golgi. Then, we removed **1** from the culture medium and replaced it with fresh medium containing **2**. We monitored fluorescence in both green and orange channels over time. Compared to control cells (washout only), the green fluorescence from **1** decayed significantly faster in cells subsequently treated with **2**, while the orange fluorescence from **2** simultaneously increased (**Figure 2F, Figure S12**). This outcome arises because the peptide-SH generated from **2** competes with the residual peptide-SH from **1** for the same pool of zDHHC enzymes needed for palmitoylation and Golgi accumulation. This competition effectively reduces the re-palmitoylation rate of peptide-SH species of **1**, accelerating the loss of its Golgi signal. These complementary experiments, one using specific enzyme inhibitors and the other using substrate competition, together demonstrate that BODIPY-CyMA relies on continuous cycles of palmitoylation and depalmitoylation for its dynamic localization at the Golgi.

These data (**Figure 2C-2F**) provide direct evidence for the existence of a dynamic steady-state, addressing the critical question of how the futile cycle enables ultrasensitive detection. Rather than functioning as a simple “back-and-forth” process, the cycle serves as a mechanism to enrich small molecules within the system. In a single-enzyme system, enzymatic assemblies are predominantly kinetically controlled, meaning that molecules diffusing away after reaction no longer participate in assembly formation. In contrast, within the enzymatic futile cycle, CyMA intermediates that diffuse, either in the Peptide-SH or Peptide-SPalm form, are continuously reprocessed and recycled back into the assembly pathway. This cycle-driven “trap” effectively shifts the system from kinetic control to a thermodynamically governed steady state, allowing the probe to reach a high local concentration even when the external precursor concentration remains at sub-nanomolar levels.

### BODIPY-CyMA generates innocuous solid-like assemblies at the Golgi

To verify that BODIPY-CyMA forms assemblies at the Golgi, we synthesized a CyMA derivative conjugated to Universal Black 2 Quencher (UBQ2, also known as Black Hole Quencher-2 or BHQ2), referred to as UBQ2-CyMA (**7**). UBQ2-CyMA incorporates a polyaromatic-azo backbone, serving as an efficient fluorescence resonance energy transfer (FRET) acceptor^[21]^ for the BODIPY fluorophore in **2**. As shown in Figure 3A, a decrease in the Golgi fluorescence intensity of **2** upon co-assembly with **7** would indicate that **2** and **7** are brought into close proximity (within 10 nm), thus suggesting the formation of co-assemblies. We treated cells with a mixture of **2** (10 nM) and varying concentrations of **7** (**Figure 3B, Figure S13**). Although **7** competes with **2** for the same zDHHC enzymes, thereby slowing the accumulation of **2**. This is reflected in a concentration-dependent increase in t_1/2_ (slower accumulation). However, this competition alone cannot account for the concentration-dependent decrease in maximum fluorescence intensity. The reduced signal is consistent with FRET-based quenching, indicating that **2** and **7** form co-assemblies at the Golgi. In contrast, co-treatment with **2** and unconjugated UBQ2, which cannot undergo S-palmitoylation, did not diminish Golgi fluorescence (**Figure S14**). These results collectively indicate that CyMA molecules form assemblies at the Golgi.

**Figure 3.**
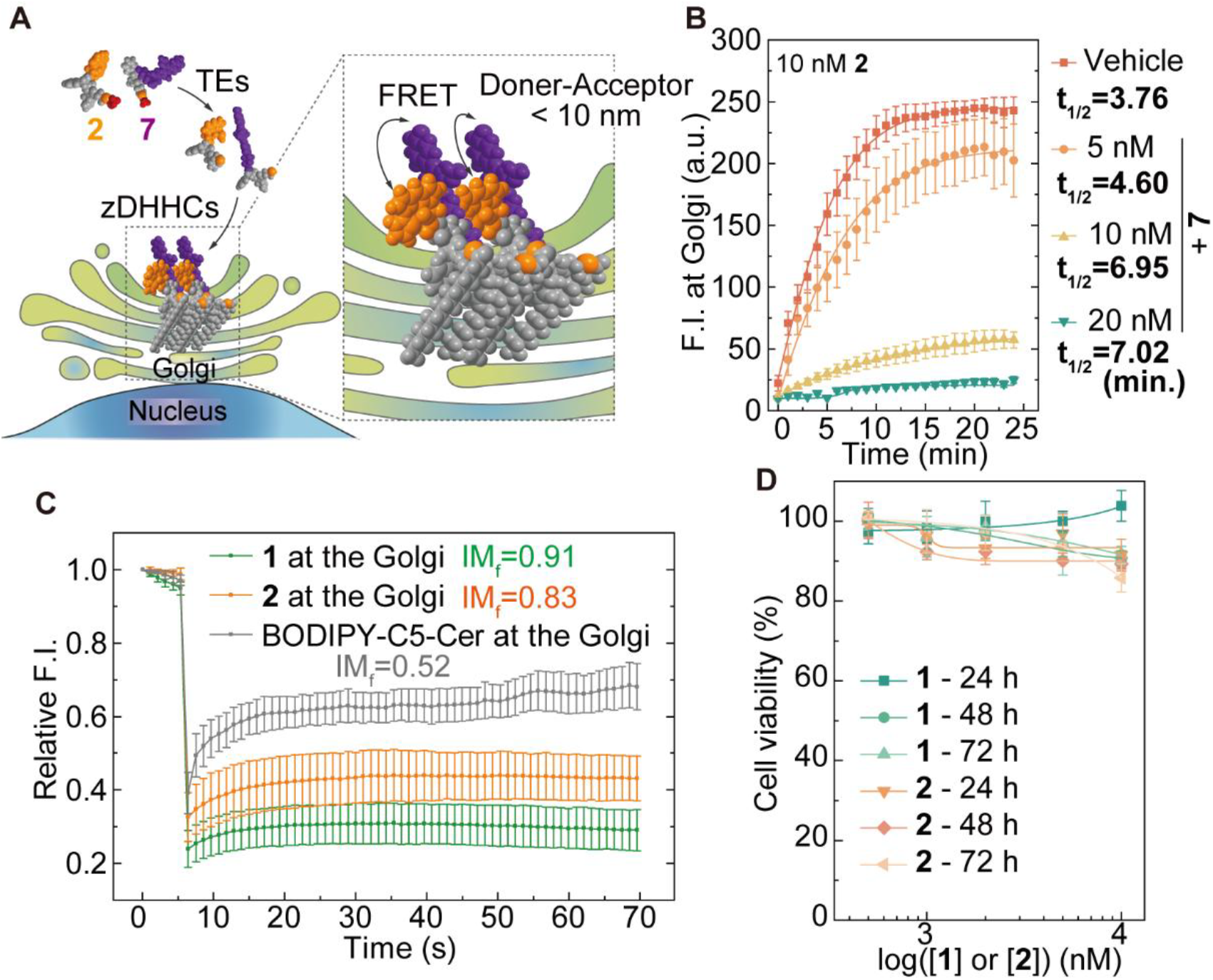
BODIPY-CyMA form assemblies at the Golgi without affecting cellular activities. (A) Schematic illustration of co-assembly of FRET doner and acceptor, **2** and **7**, respectively. (B) Golgi fluorescence intensity of HeLa cells treated with mixture of **2** (10 nM) and **7** (5, 10, 20 nM) in 24 minutes. (C) FRAP of the fluorescence from **1** (10 nM) or **2** (10 nM) or BODIPY-C5-Ceramide (500 nM) at the Golgi after 30-minute treatment. (D) Cell viability of HeLa cells treated with **1** or **2** for 24, 48, and 72 h.

To assess the impact of assembly formation on molecular mobility, we performed fluorescence recovery after photo-bleaching (FRAP) experiments. The immobile fraction (IM_f_) was determined to be 0.91 for **1** (10 nM) and 0.83 for **2** (10 nM), compared to an IM_f_ of 0.52 for the control probe BODIPY-C5-ceramide (500 nM) (**Figure 3C**). A higher IM_f_ suggests that the macroscopic assemblies form stable, solid-like structures that stay anchored at the Golgi, while also engaging in nonspecific binding of alkyl chains to the Golgi membrane. This observation of constant nanostructure is not inconsistent with the dynamic cycling of individual molecules. We propose a model where the bulk assembly serves as a stable scaffold, while individual probe molecules can dynamically exchange with soluble pool via palmitoylation/depalmitoylation cycles. Importantly, these stable assemblies were well-tolerated by cells, as continuous exposure to 10 µM of **1** or **2** over three days did not induce significant cytotoxicity (**Figure 3D**). Together, these results establish that BODIPY-CyMA form stable, non-diffusive assemblies at the Golgi without perturbing cellular viability, highlighting its utility as a robust probe for Golgi imaging.

### BODIPY-CyMA enables real-time visualization of Golgi dynamics

To demonstrate the utility of the probes, we first prelabeled cells with **2** to mark the Golgi and then treated them with BFA, a known disruptor of Golgi organization^[2a]^. As shown in **Figure 4A**, the Golgi initially appears as a distinct perinuclear structure at time zero. By 5.5 minutes post-BFA treatment, the Golgi begins to lose its defined morphology, and by 6.5 minutes, it collapses into the ER, consistent with the known mechanism of BFA-induced Golgi disassembly^[22]^. Notably, BODIPY-CyMA also reveals cell-to-cell heterogeneity in the temporal response to BFA (**Figure 4B**). For example, two cells (indicated by white arrows) display Golgi blurring at 5.5 minutes, while a third cell (blue arrow) shows similar changes at 6.5 minutes. Additional cells (green and magenta arrows) exhibit delayed responses at 7.5 and 8 minutes, respectively. These observations highlight the ability of BODIPY-CyMA to resolve dynamic and asynchronous Golgi remodeling events at single-cell resolution.

**Figure 4.**
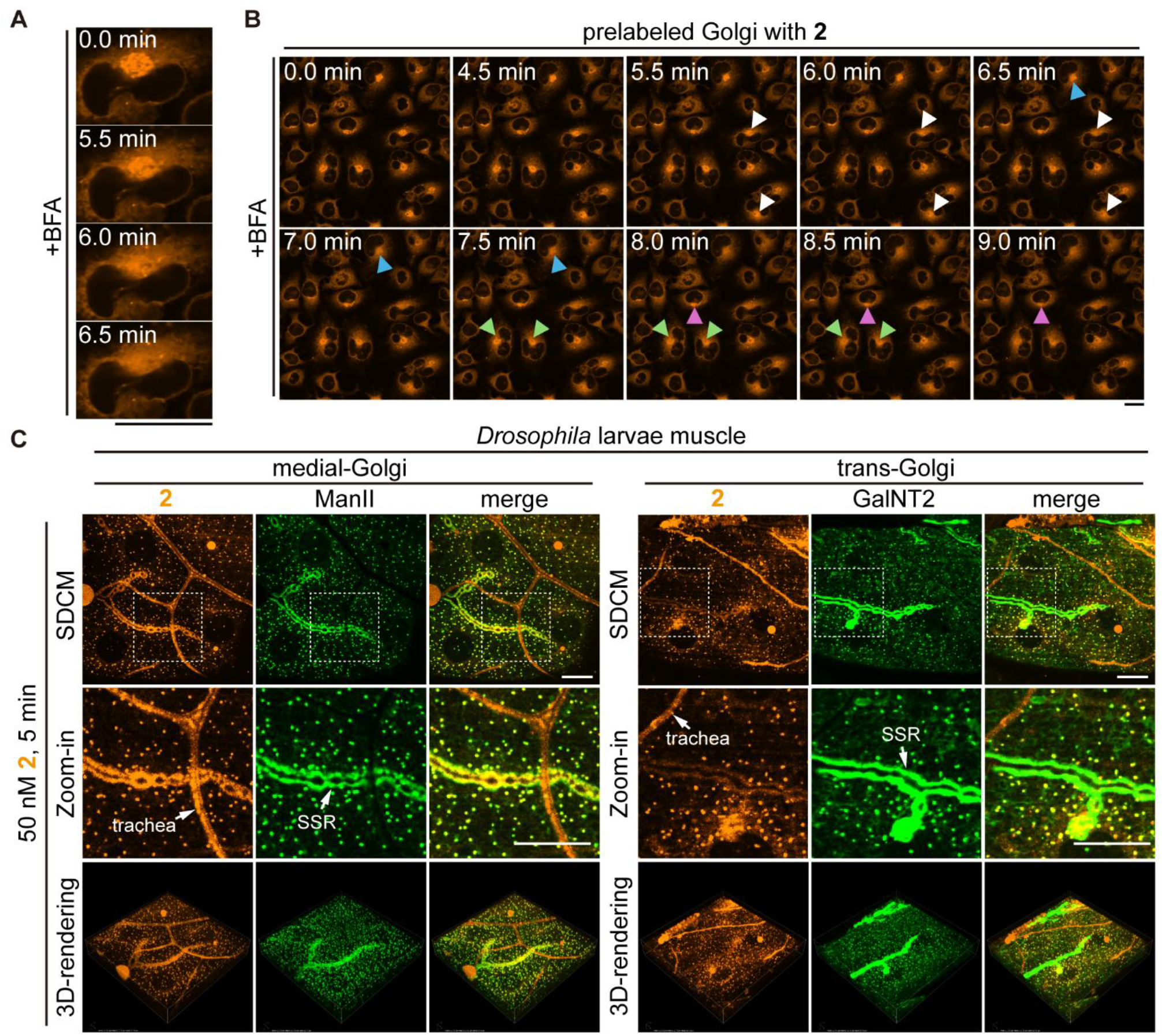
Visualization of Golgi dynamics in cultured cells and *in vivo* Golgi labeling using BODIPY-CyMA. (A) Disassembly of the Golgi in HeLa cells prelabeled with **2** (10 nM, 30 min) following treatment with BFA (10 μM). (B) Cell-to-cell heterogeneity in the Golgi response to BFA (10 μM) in HeLa cells prelabeled with **2** (10 nM, 30 min). (C) Spinning disk confocal microscopy (SDCM) and 3D-rendering of *Drosophila* larval muscle treated with **2** (50 nM, 5 min), with Golgi structures labeled using specific markers, ManII-eGFP (medial-Golgi) or GalNT2-YFP (trans-Golgi). Scale bar = 20 μm.

Further demonstrating its versatility, we used BODIPY-CyMA to visualize Golgi ministacks formed upon microtubule depolymerization with nocodazole, which fragments the Golgi into dispersed elements throughout the cytoplasm^[10]^. In nocodazole-treated cells, BODIPY-CyMA successfully labeled these ministacks (**Figure S15**), due to the continued localization of Golgi-resident enzymes (i.e., palmitoyl acyltransferases) within these structures^[23]^. Importantly, Golgi fluorescence was lost upon cell fixation, underscoring the requirement for active, *in situ* palmitoylation–depalmitoylation cycles in maintaining Golgi targeting (**Figure S16**).

### BODIPY-CyMA for *in vivo* Golgi imaging

To evaluate its performance beyond cultured cells, we examined its Golgi-labeling capability in *Drosophila* larval muscle as an *in vivo* model (**Figure 4C**). Remarkably, treatment with 50 nM of **2** resulted in rapid and efficient Golgi labeling within 10 minutes, as indicated by strong colocalization of the orange fluorescence from BODIPY-CyMA with green fluorescent markers for the medial-Golgi (ManII) and trans-Golgi (GalNT2), expressed using the GAL4-UAS system with a muscle-specific driver. In contrast, although BODIPY-CyMA also labeled Golgi structures in the trachea, no green fluorescence was observed in this tissue due to the absence of Golgi marker expression outside of muscle cells. These results highlight the high Golgi-targeting efficiency and practical utility of BODIPY-CyMA *in vivo*.

Additionally, BODIPY-CyMA accumulated in the subsynaptic reticulum (SSR), a membrane-rich structure that connects synaptic contacts to the underlying muscle^[24]^. Interestingly, minimal fluorescence was detected in the motor neuron presynaptic terminals contacting the SSR (**Figure S17**), and extended incubation following a pulse of BODIPY-CyMA did not enhance neuronal labeling (**Figure S18**), consistent with the predicted absence of Golgi from presynaptic terminals.^[25]^ Together, these findings demonstrate that BODIPY-CyMA is a robust probe for *in vivo* visualization of the Golgi.

## Conclusion

This work introduces BODIPY-CyMA, an imaging probe that converts an endogenous enzymatic futile cycle into a self-sustaining molecular assembly for ultrasensitive and dynamic Golgi imaging. The work demonstrates how the rational engineering of a CyMA scaffold that harnesses an endogenous enzymatic futile cycle involving S-palmitoylation by zDHHC enzymes and depalmitoylation by TEs. This cycle mediates the selective accumulation and retention of BODIPY-CyMA at the Golgi. Importantly, the probe undergoes *in situ* activation and localization through enzymatic processing, ensuring that fluorescent signals are only generated at the site of interest within the native cellular context. The superior photophysical properties of the BODIPY fluorophore, including high quantum yield, minimal photobleaching, and sharp emission profiles, enable sensitive detection at picomolar concentrations and clean spectral separation from commonly used fluorescent proteins (e.g., GFP, YFP, mCherry), allowing for multiplexed imaging in complex biological environments. Operating at ultralow concentrations minimizes artifacts associated with high concentrations of lipophilic dyes^[26]^ and allowing accurate, non-perturbative observation of cellular processes. Together, these attributes make BODIPY-CyMA exceptionally well-suited for both live-cell and *in vivo* applications.

Given that the human cells contain at least 23 palmitoyl acyltransferases, which are present on multiple organelles including the ER and plasma membrane, it is remarkable that BODIPY-CyMA localized at Golgi. The remarkable selectivity likely arises from the CyMA futile cycle. While the factors contribute to Golgi localization has yet to be confirmed, Golgi localized TEs and zDHHCs likely play the key roles. This observation also support that Golgi is the palmitoylation site for both vesicular^[20b]^ and nonvesicular^[27]^ protein trafficking. Beyond its utility as an organelle marker, BODIPY-CyMA act as a dynamic reporter of enzymatic activities. While existing methods, such as labeling with chemical reporters via click reactions^[28]^, can identify protein modification, but they typically require endpoint measurements. In contrast, the accumulation of BODIPY-CyMA is controlled by an enzymatic switch, which is the balance between the counteracting activities of zDHHCs and TEs. This dual-enzyme mechanism is conceptually distinct from probes that rely on a single enzymatic trigger for signal accumulation^[29]^. Consequently, the steady-state signal intensity of the probe serves as a direct, real-time readout of the palmitoylation-depalmitoylation equilibrium at the Golgi in living cells. This capability to sense enzymatic activity from both sides is particularly valuable, as methods to evaluate protein palmitoylation dynamics in living cells remain limited^[30]^. Therefore, by leveraging a design principle established for creating functional biosensors, BODIPY-CyMA provides a novel chemical tool to visualize the integrated enzymatic activity that regulates this reversible post-translational modification.

The modularity of the CyMA platform affords spectral and functional versatility through facile fluorophore substitution. Beyond enabling advanced modalities like super-resolution imaging^[31]^, this tunability facilitates the investigation of Golgi dynamics across diverse cellular contexts, such as variations in pH^[32]^, polarity^[33]^, and membrane tension^[34]^, and supports compatibility with a range of imaging platforms, for example, GLIM^[35]^ and IR^[29c]^. This flexibility enables CyMA a versatile tool for probing not only Golgi structure, but also Golgi-associated biological processes, including protein trafficking^[36]^, lipid metabolism^[37]^, posttranslational modifications^[38]^, and mitosis^[39]^. In conclusion, by harnessing endogenous enzymatic activity for *in situ* signal amplification and retention, BODIPY-CyMA offers a promising addition to the imaging toolkit for studying Golgi dynamics with minimal perturbation.

## Supporting Information

Materials and methods, Figures S1 to S21 (PDF)

Time-series CLSM of HeLa cells treated with **1** or **2** (10 nM) over 30 minutes. (AVI)

## Acknowledgements

We thank the Brandeis Light Microscopy Facility (RRID:SCR_025892) for assistance. This work is partially sup-ported by NIH (CA142746) (BX), NSF (DMR-2011846) (BX) and grant 2023-321163 from the Chan Zuckerberg Initiative Donor-Advised Fund at the Silicon Valley Community Foundation (BX), NIH NS103967 (AR).

