## Supplementary information for "Cycling Molecular Assemblies (CyMA) for Ultrasensitive Golgi Imaging"

**This PDF file includes:**

Materials and methods

Figures S1 to S21

Legends for Movies S1

**Other supporting materials for this manuscript include the following:**

Movie S1

**Materials and Methods**

**Materials**

2-Cl-trityl chloride resin (1.0 mmol/g), Fmoc protected amino acid, and HBTU were obtained from GL Biochem (Shanghai, China). N, N-diisopropylethylamine (DIEA), 2-Mercaptoethanol, chloroquine diphosphate salt, and solvents were obtained from Fisher Scientific. Acetyl chloride were purchased from TCI America. DC661 was purchased from MedChemExpress. ML-211 was purchased from APExBIO. 2-bromohexadecanoic acid was purchased from Sigma-Aldrich. All the chemical reagents and solvents were used as received from commercial sources without further purification.

Xfect™ Transfection Reagent (Catalog # 631317) was purchased from TaKaRa. mEmerald-ELP-1-25 was a gift from Michael Davidson (Addgene plasmid # 54080; http://n2t.net/addgene:54080; RRID:Addgene_54080). mNeonGreen-Giantin was a gift from Dorus Gadella (Addgene plasmid # 98880; http://n2t.net/addgene:98880; RRID:Addgene_98880). EGFP-GalT was a gift from Jennifer Lippincott-Schwartz (Addgene plasmid # 11929; http://n2t.net/addgene:11929; RRID:Addgene_11929). mEmerald-TGNP-N-10 was a gift from Michael Davidson (Addgene plasmid # 54279; http://n2t.net/addgene:54279; RRID:Addgene_54279).

Minimum Essential Medium (MEM), Dulbecco's Modified Eagle Medium (DMEM), McCoy's 5A, fetal bovine serum (FBS) Opti-MEM, CO_2_ Independent Medium and penicillin-streptomycin (PS) were purchased from Gibco. RPMI 1640 medium, F-12K medium, and Eagle's Minimum Essential Medium (EMEM) were purchased from American Type Culture Collection (ATCC, USA).

**Instruments**

All precursors and compounds were purified by a reverse phase HPLC (Agilent 1100 Series) equipped with an XTerra C18 RP column. HPLC grade acetonitrile (0.1% TFA) and HPLC grade water (0.1% TFA) were used as the eluents. LC-MS spectra were obtained with a Waters Acquity Ultra Performance LC with Waters MICROMASS detector and a Bruker Elute PLUS UHPLC with a Bruker timsTOF Pro. Fluorescence microscopes are described in detail below.

**Methods**

**Synthesis of** **Golgi targeting backbone**

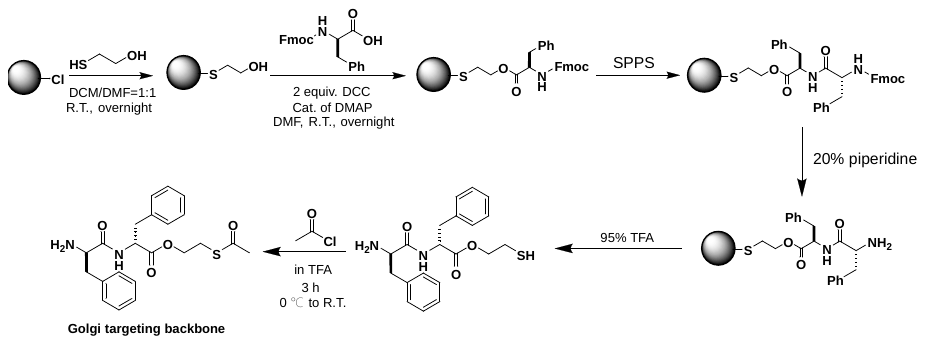

**Scheme S1.** Synthetic procedure of Golgi targeting backbone.

As shown in **Scheme S1**, we used standard Fmoc chemistry for solid phase peptide synthesis (SPPS) with 2-chlorotrityl chloride resin and Fmoc-protected amino acids with appropriately protected side chains. Briefly, the 2-Cl resin (1 g) was swelling in dry DCM for 30 minutes, followed by loading the first amino acid onto the resin. Next, 10 equivalents of 2-mercaptoethanol dissolved in DCM/DMF (v:v=1:1) were added into the SPPS reactor and incubated with the resin overnight at room temperature. The thiol group of 2-mercaptoethanol is selectively attached to the resin due to its high nucleophilicity. After discarding the liquid phase, the resin with mercaptoethanol linker was washed five times with DMF. Subsequently, Fmoc-protected amino acid (1.5 equiv.), along with N,N′-dicyclohexylcarbodiimide (DCC, 2 equiv.) and a catalytic amount of 4-Dimethylaminopyridine (DMAP) dissolved in DMF, was added to the reactor and incubated with the resin overnight at room temperature. The following day, the Fmoc group was removed with 20% piperidine in DMF, and the next Fmoc-protected amino acid was coupled to the free amino group using HBTU as the coupling reagent. After removal of the Fmoc protecting group, the peptide chain was cleaved from the resin by 95% TFA in water for 1 hour. The resulting peptide with thiol handle is directly reacted with an acyl chloride in TFA to generate a thioester in high yield. The crude Golgi targeting backbone (ffesSAc) are further purified by reverse-phase HPLC to yield the final product.

**Synthesis of BODIPY-CyMA precursor (Compound 1, 2, 7)**

The purified Golgi-targeting backbone (ffesSAc, 1 equiv.) was dissolved in anhydrous DMF, and the pH of the solution was adjusted to approximately 8 using DIPEA. Subsequently, BODIPY-NHS or UBQ2-NHS (1.1 equiv.) was added, and the reaction mixture was stirred for 4 hours at room temperature. Upon completion, the mixture was purified by reverse-phase HPLC to afford the final product.

**Cell culture**

HeLa cells were purchased from ATCC. Cell line was authenticated by CellCheck 9 - human (9 Marker STR Profile and Inter-species Contamination Test, IDEXX), confirming 100% match of the cell identity. HeLa cells were cultured in MEM supplemented with 10% FBS. All the cell lines were supplemented with 100 U/mL penicillin and 100 µg/mL streptomycin and were cultured and humidified with 5% CO_2_ at 37 °C.

**Cell transfection**

Cells were seeded at 1.5×10^5^ cells per confocal dish for 24 h to allow attachment. Upon reaching 40-50% confluency, the cells were transfected with Xfect™ Transfection Reagent. Specifically, 5 µg of the plasmid DNA was diluted with Xfect Reaction Buffer, followed by the addition of 1.5 μL Xfect Polymer. The mixture was briefly vortexed and incubated for 10 min at room temperature. The entire 100 μL of nanoparticle complex solution was then added dropwise to the cell culture medium, and the dish was rocked briefly. The confocal dish was incubated at 37°C overnight, and the media was replaced with fresh culture media for an additional 48-hour incubation. The cells were then ready for live-cell imaging.

**Confocal microscopy**

A confocal dish (35 mm dish with 20 mm bottom well, #1.5 glass) was used to prepare CLSM samples. For live-cell imaging, cells in exponential growth phase were seeded on the confocal dish at 1.0×10^5^ cells per dish and incubated for 24h. After removing the culture medium, fresh medium containing the compound of interest was added to the cells for co-incubation at 37 °C in a humidified atmosphere of 5% CO_2_ for the desired period. Afterwards, the nuclei of cells were stained with Hoechst 33342 for 10 minutes, and the samples were washed with 1 mL of Live Cell Imaging Solution four times to fully remove the residual Hoechst 33342.

For time-lapse live-cell imaging, cells in exponential growth phase were seeded on a confocal dish at 1.0×10^5^ cells per dish and incubated for 24 h. The samples were washed with 1 mL of Live Cell Imaging Solution three times, and the nuclei were stained with Hoechst 33342 for 10 minutes. The samples were then washed with 1 mL of Live Cell Imaging Solution four times to remove the residual Hoechst 33342. Cells were imaged on a Nikon AX-R resonant confocal system using a 60×/1.4 Oil objective. The position of cells and the focal plane of the laser beam were determined using the fluorescence from the stained nuclei with a laser with 405 nm wavelength, and the Nikon Perfect Focus System was activated to prevent focus drift. The imaging solution in the confocal dish was replaced with fresh imaging solution containing the compound of interest. CLSM images of different channels were then recorded, with the time-series interval set to be 1 minute or no-delay. After a specified number of imaging cycles, the fluorescence images from both channels were saved for further analysis.

**Cell pretreated with inhibitors**

HeLa cells (1.5×10^5^ cells) were seeded in a confocal dish for 24 h to allow attachment. The culture media was then replaced with fresh medium containing inhibitors for LYPLA1/2 (ML211, 50 μM), PPT1 (DC661, 20 μM), palmitoylacyltransferases (2-BP, 10 μM), and the cells were incubated for 30 minutes at 37 °C. Afterward, the cell medium was replaced with fresh medium containing BODIPY-CyMA for live-cell imaging.

**FRAP assay**

FRAP was performed on a Zeiss LSM 880 confocal microscopy using a 63×/1.4 Oil objective or a Nikon AX-R resonant confocal system using a 60×/1.4 Oil objective. Five pre-bleach images were captured, followed by photobleaching using 488-nm laser at 100% intensity within a selected region. Another region was imaged without photobleaching as an internal control. 512×512-pixel images were captured at 0.26 s (or 5.02 s for Nikon AX-R CLSM) intervals using a 488-nm laser at 100% intensity with the pinhole set at 1 airy unit. Imaging continued until no further recovery was observed. The fluorescence recovery of the photobleached region was normalized and fitted into an exponential function.

**Palmitoylation of BODIPY-CyMA characterized by LC/HR-MS**

A total of 2.4×10^7^ cells were treated with 500 nM of BODIPY-CyMA or vehicle (DMSO) for 30 minutes, and then washed with HEPES buffer twice. Cells were collected and centrifuged to obtain a pellet. The pellet was resuspended in 500 µL of HEPES buffer, followed by the addition of 1.5 mL of DCM and 2 mL of methanol. The mixture was allowed to sit for 10 minutes. Afterward, 0.5 mL DCM and 0.5 mL Tris-HCl (50 mM, pH 2.0) were added, and the tube was centrifuged to collect the organic phase. The organic phase was washed with buffer (2 mL methanol+1 mL Tris-HCl (50 mM, pH 2.0)) and centrifuged again to obtain the organic phase. The organic solvent was evaporated with N_2_, and the remaining solid was dissolved in 100 µL of methanol and sent for LC/HR-MS analysis.

***Drosophila* culture and imaging**

Flies were cultured using standard media and techniques and maintained at 25°C. The following strains were used and obtained from the Bloomington Drosophila Stock Center: UAS-ManII-eGFP (RRID:BDSC_65248) and UAS-GalNT2-YFP (RRID:BDSC_65252 For muscle expression C57-Gal4 (PMID 8893021) was used. For imaging of larval muscle, wandering third instar larvae were pinned down and filleted one at a time in HL3.1 on slides containing sylgard in a silicone mold. The larvae were treated for 10 minutes in HL3.1 containing 50 nM **2**. Following treatment, the larvae were quickly washed 2 times in HL3.1 then 50 µL of HL3.1 was added to the larva and covered with a coverslip for imaging and imaged as soon as possible (< 5min). Z-stacks were acquired using a Nikon Ni-E upright microscope equipped with a Yokogawa CSU-W1 spinning disk head, an Andor iXon 897U EMCCD camera, and Nikon Elements AR software. A 60X (n.a. 1.4) oil immersion objective was used to image larval body wall muscle 4.

**MTT assay**

The MTT assay was used to determine cell viability for cytotoxicity evaluation. Cells were seeded at 1×10^4^ cells per well in 96-well plates for 24 h to allow attachment. Culture media were replaced with fresh culture media containing the compounds at a series of concentrations. After 24, 48, and 72 hours, 10 µL of MTT solution (5 mg/mL) was added to each well, and the plate was incubated in the dark for 4 h at 37 °C. Then, 100 µL of 10% SDS-HCl was added to stop the reaction and dissolve the formazan. The absorbance at 595 nm was determined by a microplate reader. The assay was repeated three times, and the mean values of three measurements were plotted, with error bars representing standard deviation. For 7-day cytotoxicity, seed cells at 5,000 cells per well, and fresh D-peptide-containing medium was added every 3 days.

**Quantification and Data Analysis**

**All graphs were created using Origin 2021. Quantification of fluorescence intensity was conducted using Fiji. Error bars represent s.d. unless otherwise noted. For comparison between two groups, *p* values were determined using two-tailed Student’s t-tests. All cell-based or cell-free experiments were repeated two to three times.**

**Supplementary figures**

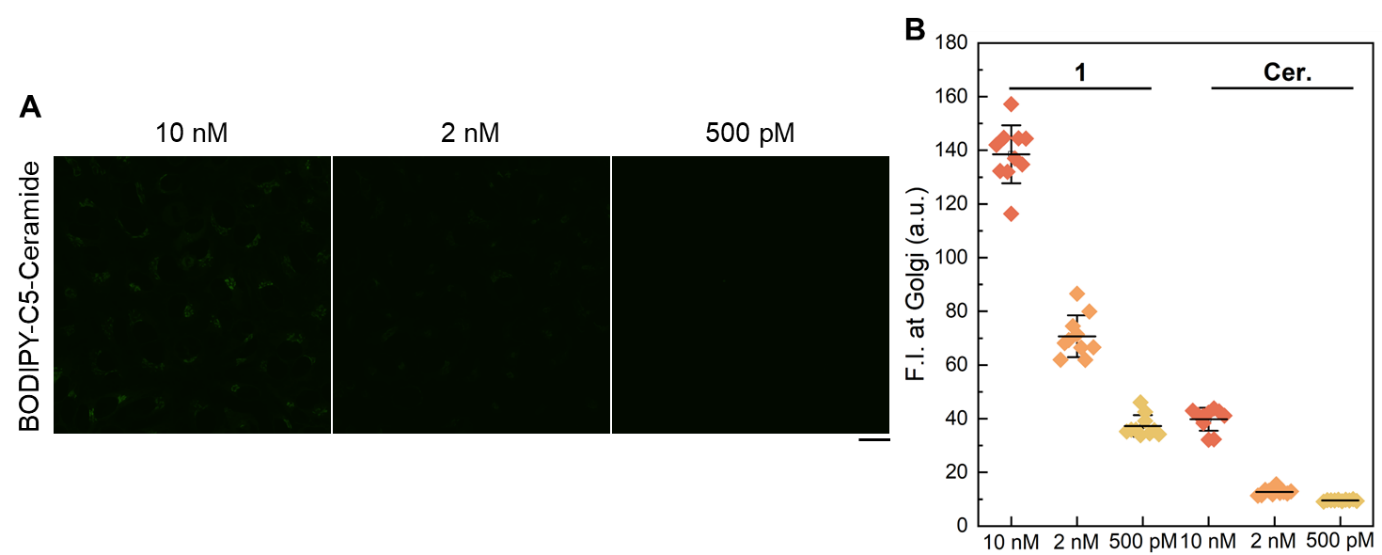

**Figure S1.** (A) CLSM of HeLa cells treated with BODIPY-C5-ceramide (10 nM, 2 nM, 500 pM) for 4 hours. (B) Quantitative Golgi fluorescence intensity of HeLa cells treated with BODIPY-CyMA (**1**) or BODIPY-C5-ceramide (n = 10). Scale bar = 20 μm.

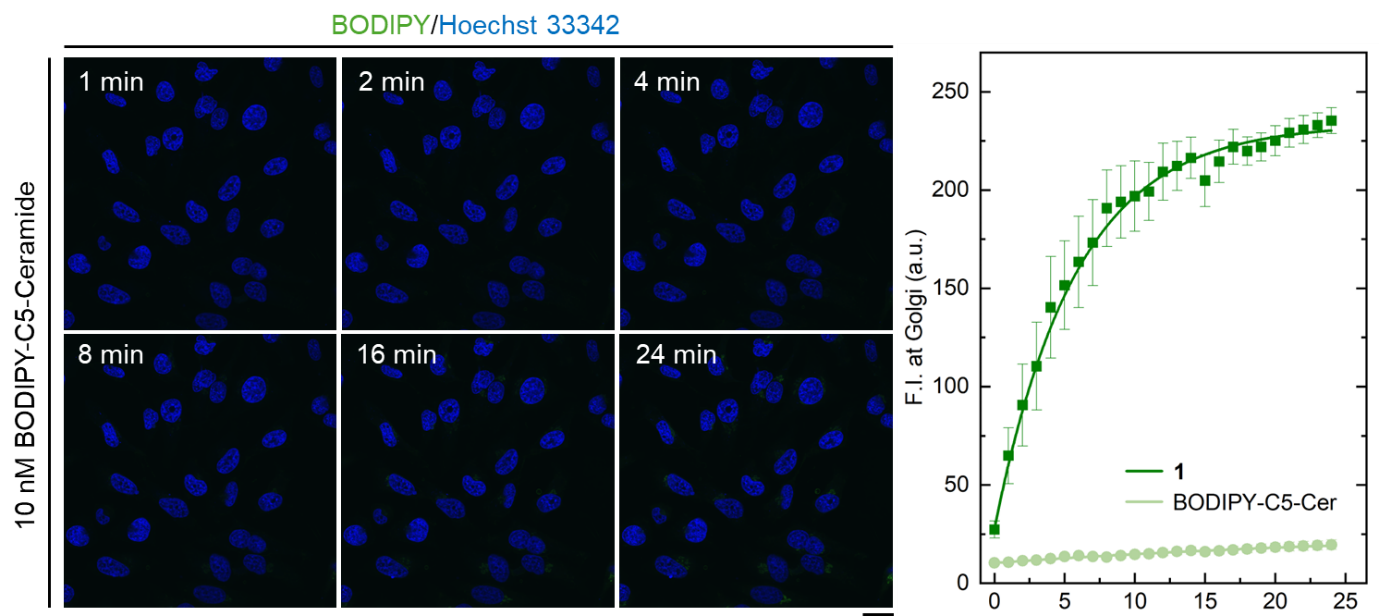

**Figure S2.** (A) CLSM of HeLa cells treated with BODIPY-C5-ceramide (10 nM) over 24 minutes. (B) Quantitative Golgi fluorescence intensity of HeLa cells treated with BODIPY-CyMA (**1**, 10 nM) or BODIPY-C5-ceramide (10 nM) over 24 minutes (n = 10). Scale bar = 20 μm.

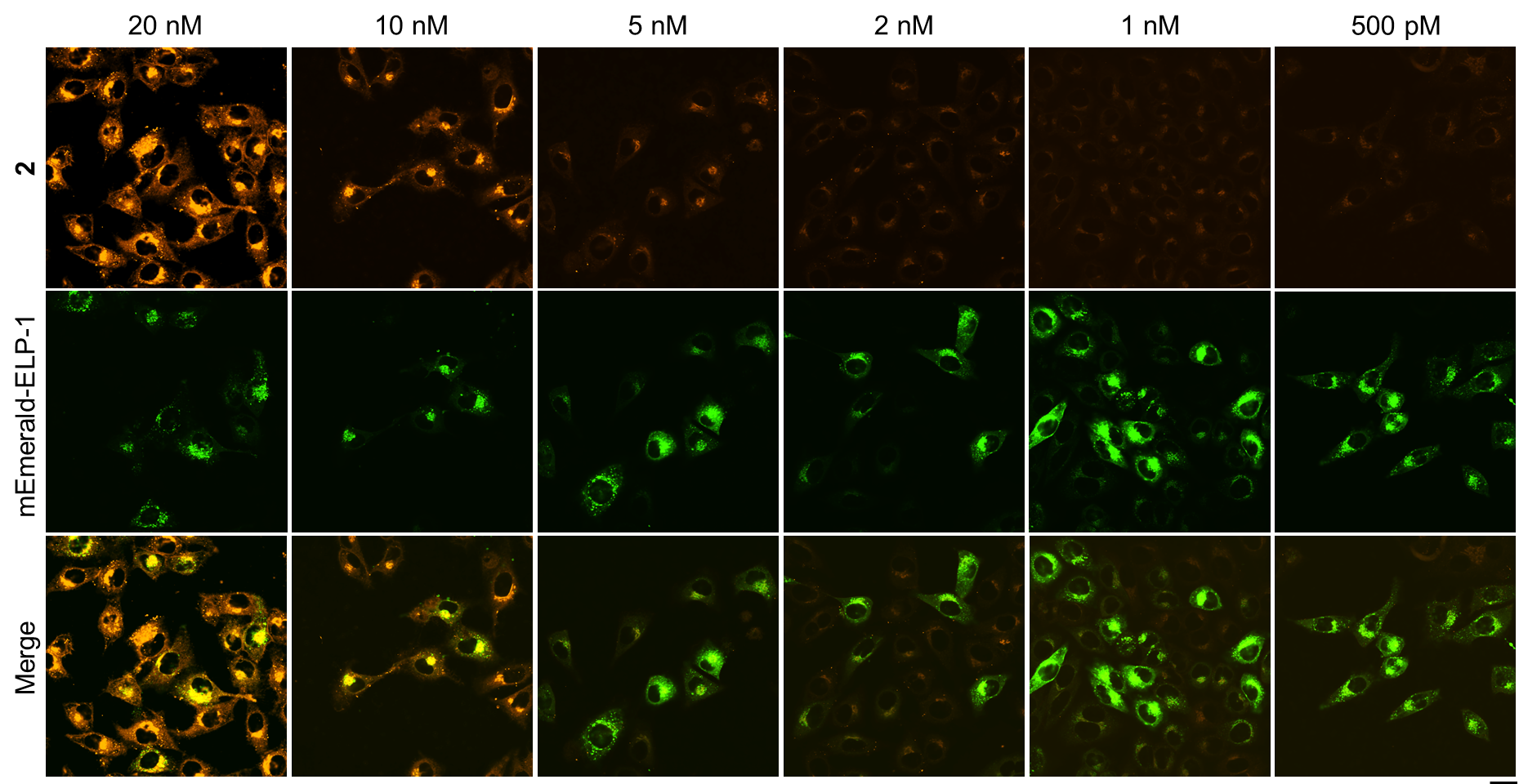

**Figure S3.** CLSM of mEmerald-ELP-1 (KDELR2) transfected HeLa cells treated with **2** (20 nM, 10 nM, 5 nM, 2 nM, 1 nM, 500 pM) for 4 hours. Scale bar = 20 μm.

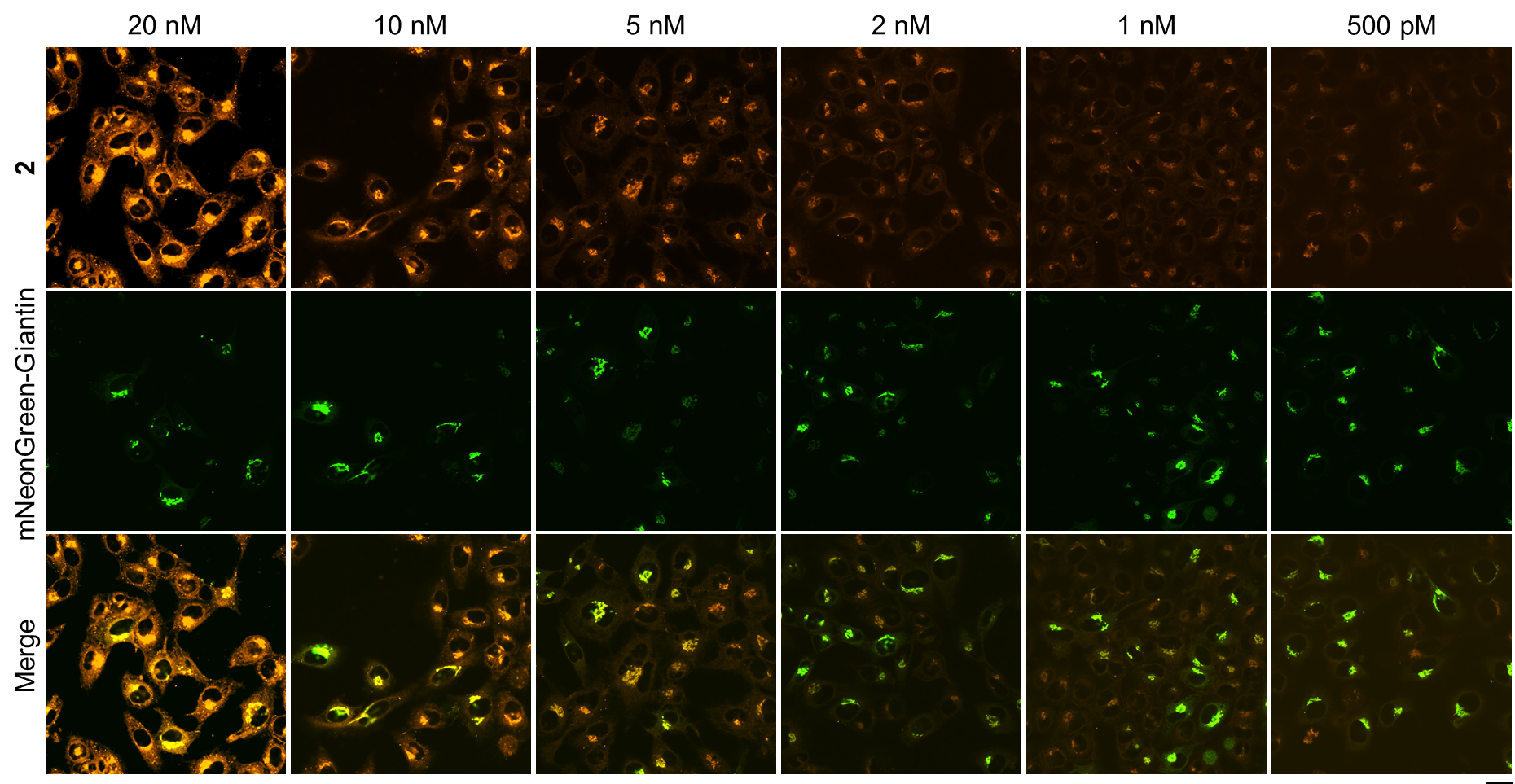

**Figure S4.** CLSM of mNeonGreen-Giantin transfected HeLa cells treated with **2** (20 nM, 10 nM, 5 nM, 2 nM, 1 nM, 500 pM) for 4 hours. Scale bar = 20 μm.

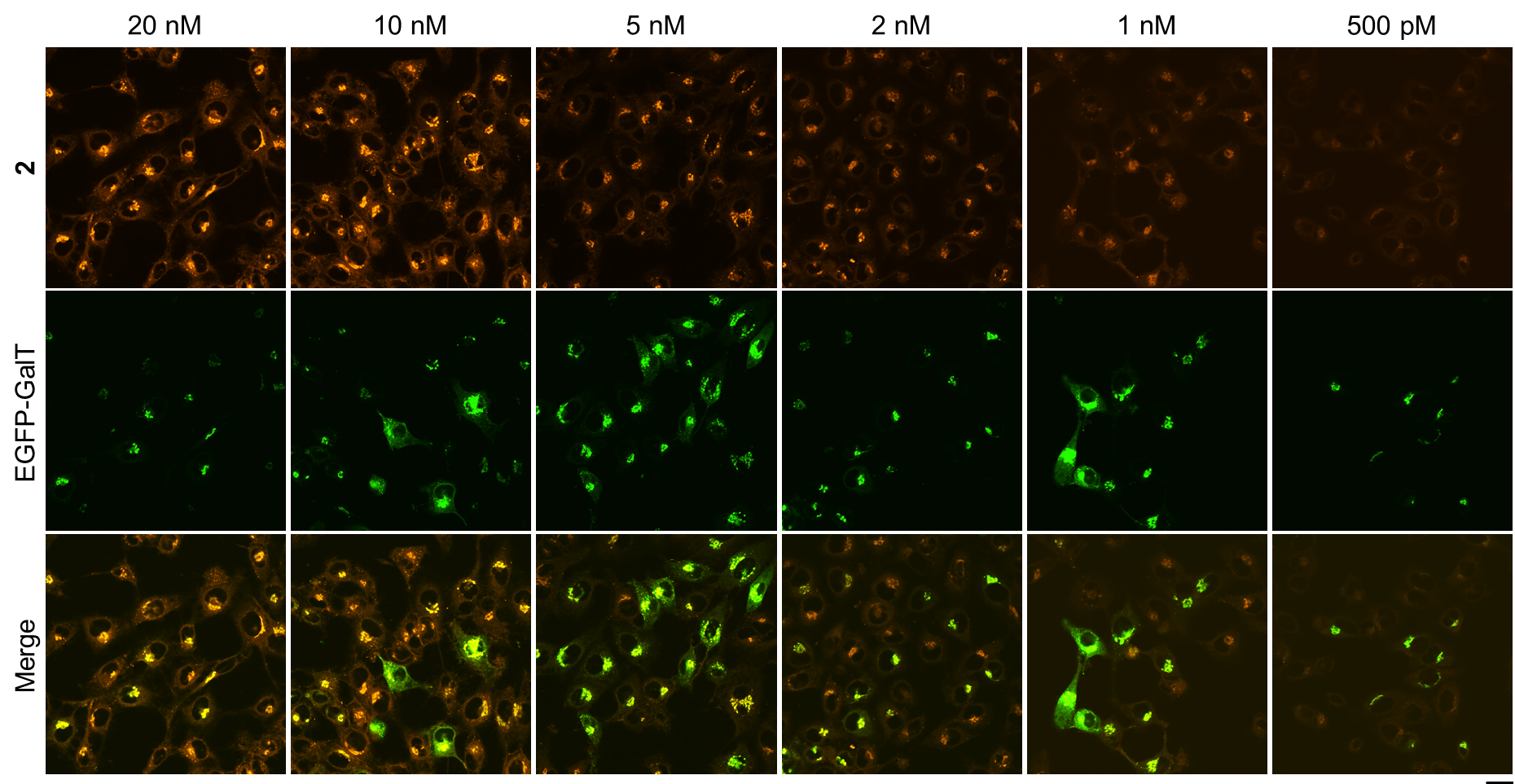

**Figure S5.** CLSM of EGFP-GalT transfected HeLa cells treated with **2** (20 nM, 10 nM, 5 nM, 2 nM, 1 nM, 500 pM) for 4 hours. Scale bar = 20 μm.

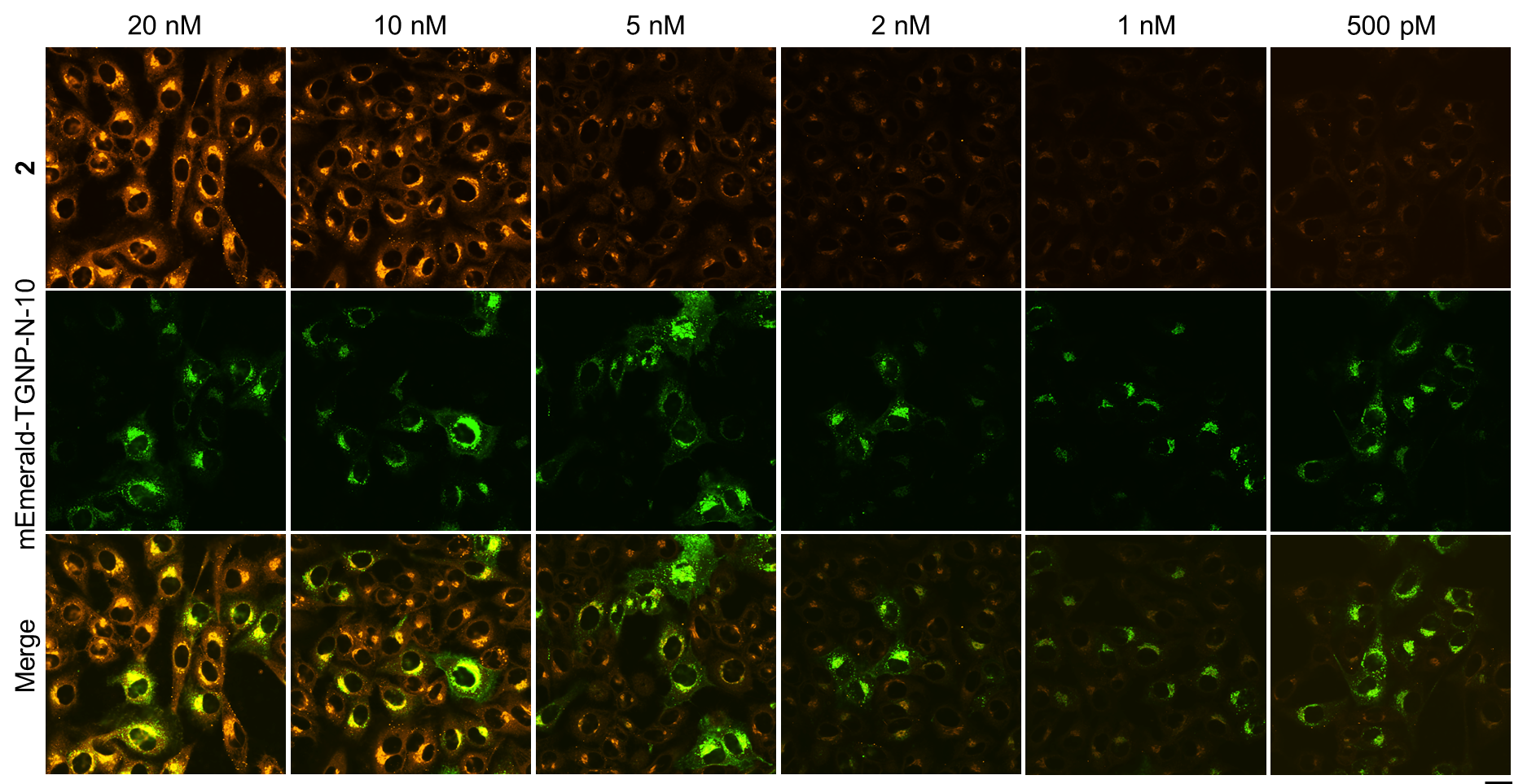

**Figure S6.** CLSM of mEmerald-TGNP-N-10 (TGN46) transfected HeLa cells treated with **2** (20 nM, 10 nM, 5 nM, 2 nM, 1 nM, 500 pM) for 4 hours. Scale bar = 20 μm.

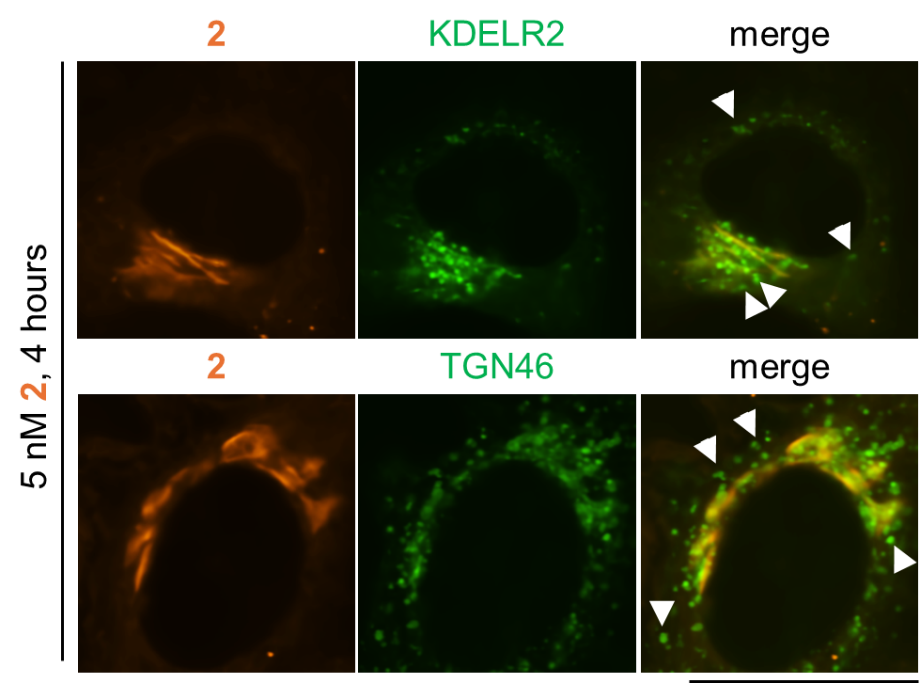

**Figure S7.** CLSM of mEmerald-ELP-1 (KDELR2) or mEmerald-TGNP-N-10 (TGN46) transfected HeLa cells treated with **2** (5 nM) for 4 hours. White arrows indicate the Golgi related transport vesicles. Scale bar = 20 μm.

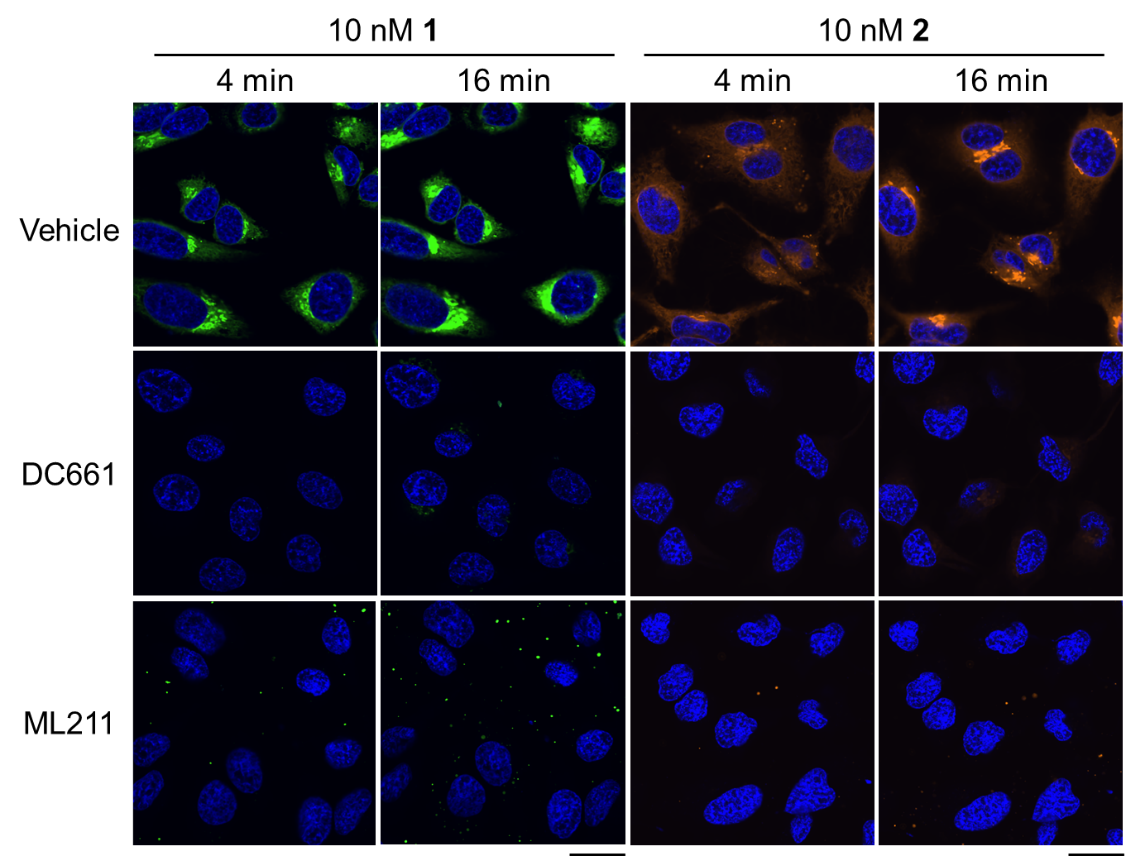

**Figure S8.** CLSM of HeLa cells pretreated with thioesterases inhibitor (DC661, 20 μM or ML211, 50 μM) for 30 minutes and then incubated with **1** or **2** (10 nM) over 16 minutes. Scale bar = 20 μm.

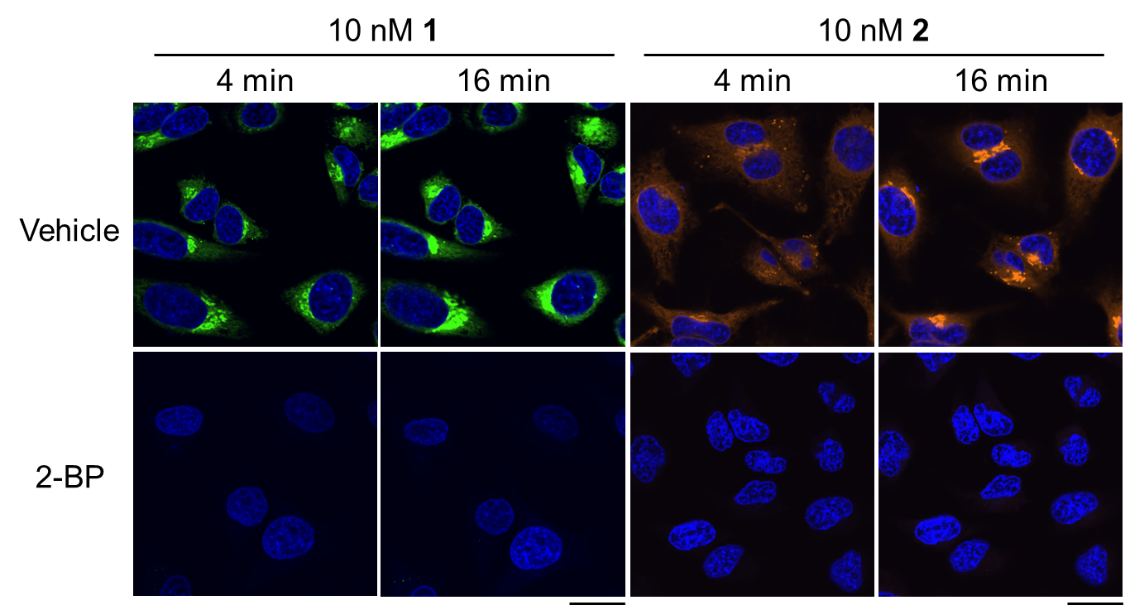

**Figure S9.** CLSM of HeLa cells pretreated with zDHHCs inhibitor (2-BP, 10 μM) for 30 minutes and then incubated with **1** or **2** (10 nM) over 16 minutes. Scale bar = 20 μm.

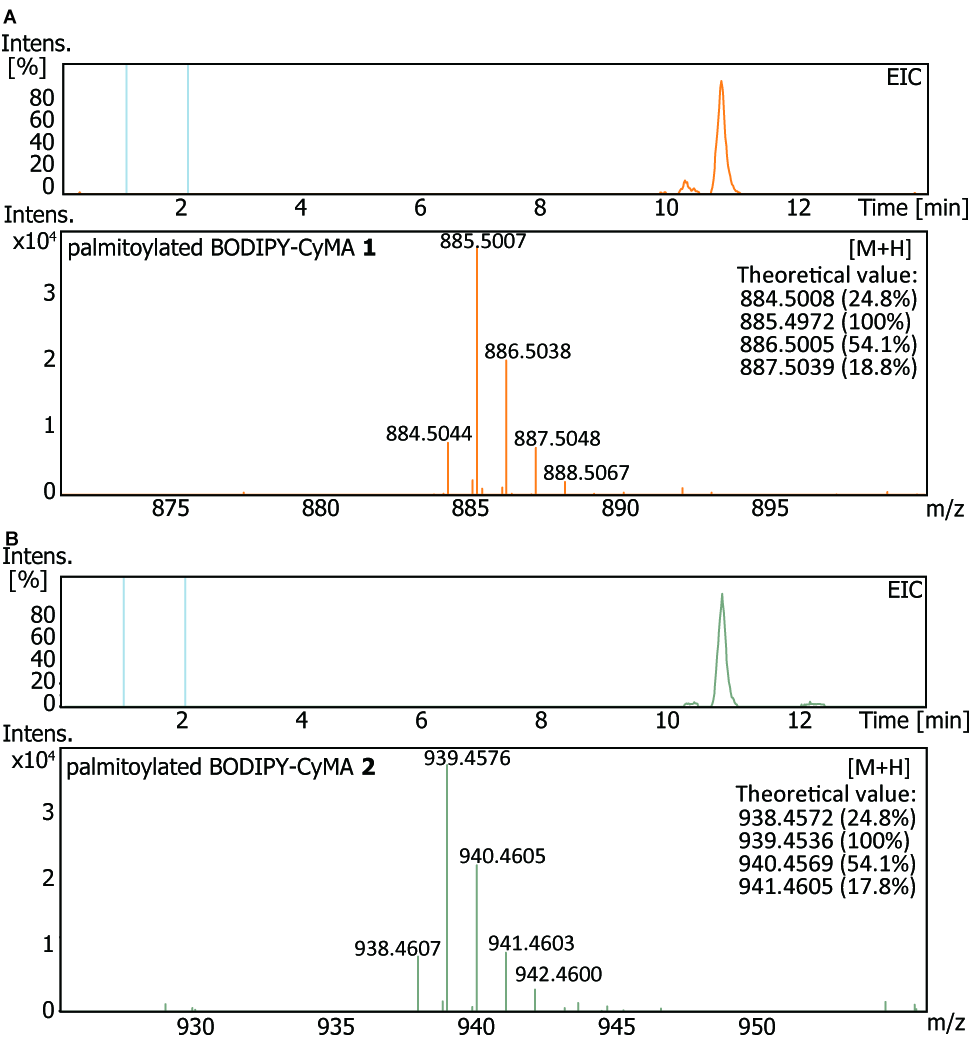

**Figure S10.** Extracted ion chromatogram **(**EIC) and HRMS of palmitoylated BODIPY-CyMA **1** (A) and **2** (B) in lysate of HeLa cells treated with **1** or **2** (500 nM) for 30 minutes.

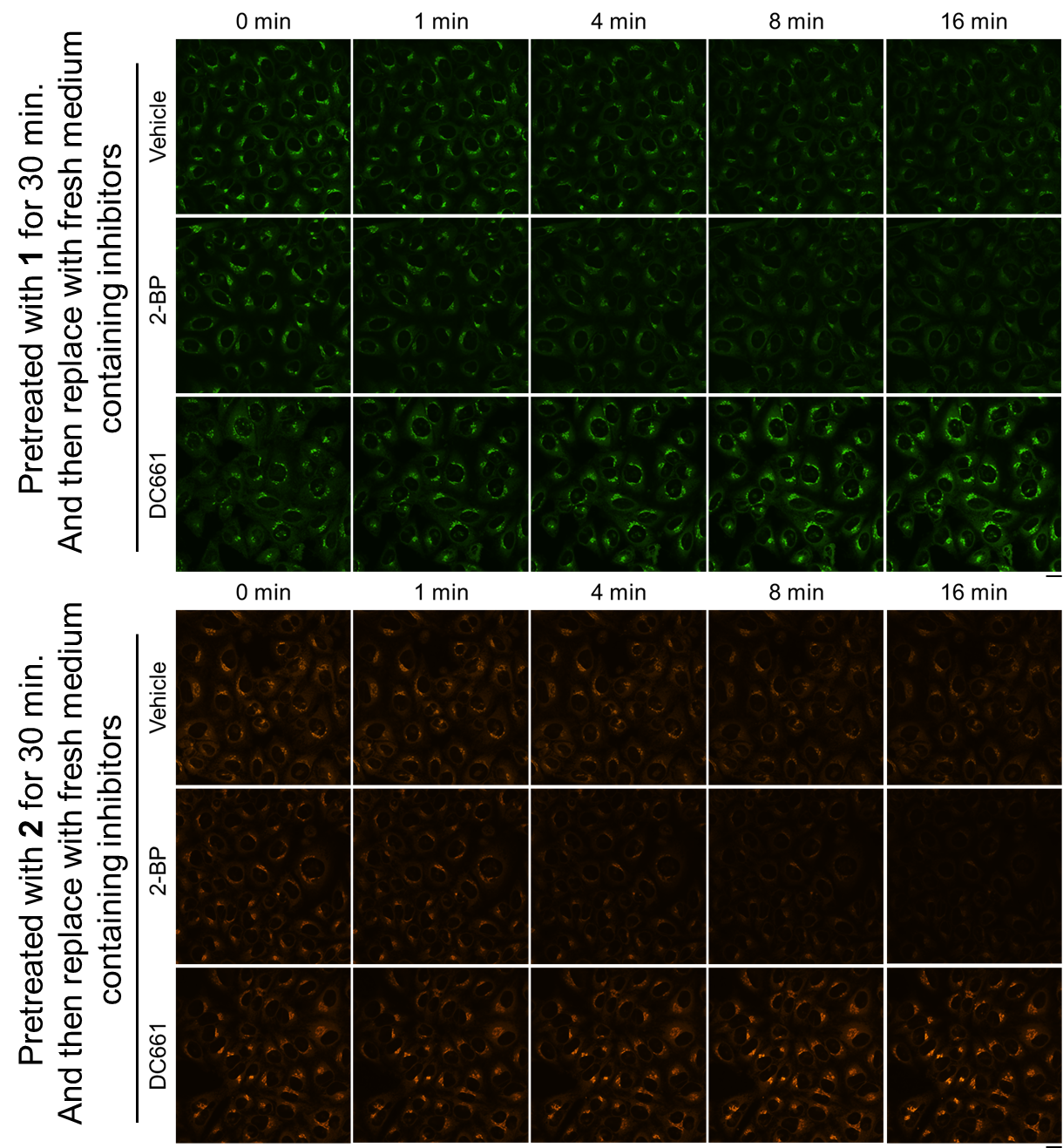

**Figure S11.** CLSM of HeLa cells pretreated with **1** or **2** (10 nM, 30 min), then switched to fresh media with vehicle or either PPT1 inhibitor (DC661, 20 μM) or zDHHC inhibitor (2-BP, 20 μM) over 16 minutes. Scale bar = 20 μm.

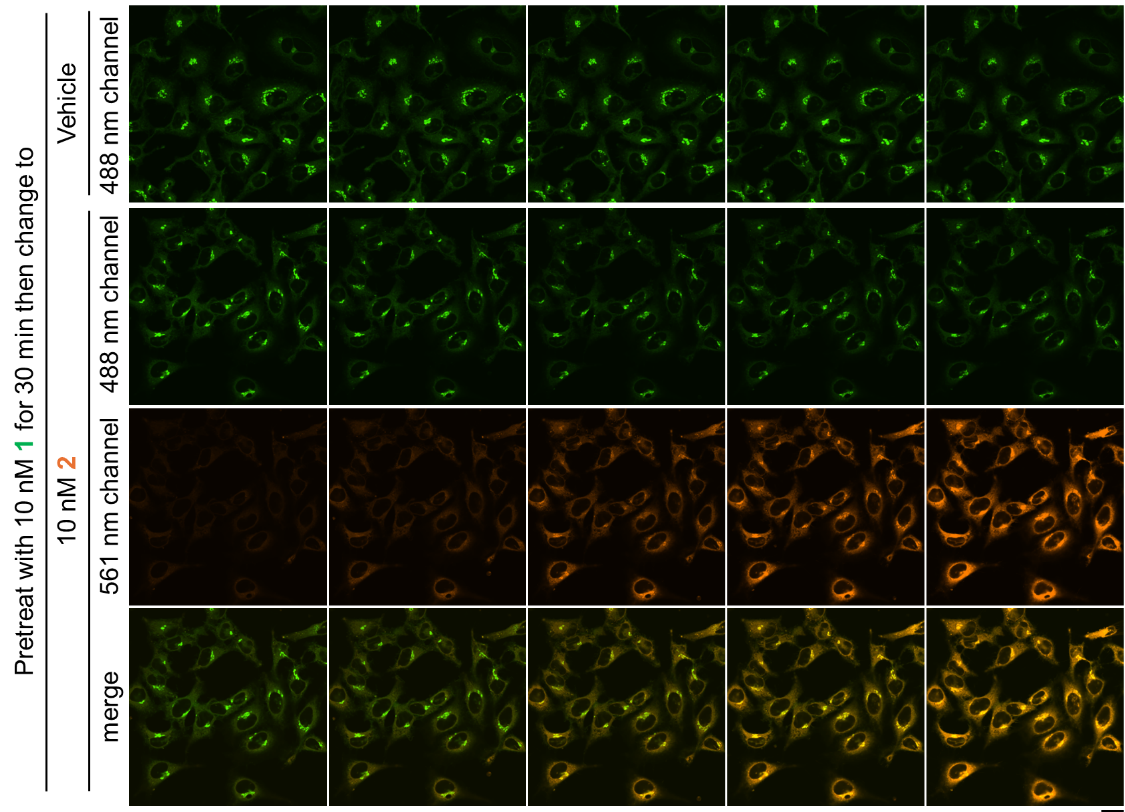

**Figure S12.** CLSM of HeLa cells were pretreated with **1** (10 nM, 30 min), then switched to fresh media with vehicle or **2** (10 nM, 30 min). Scale bar = 20 μm.

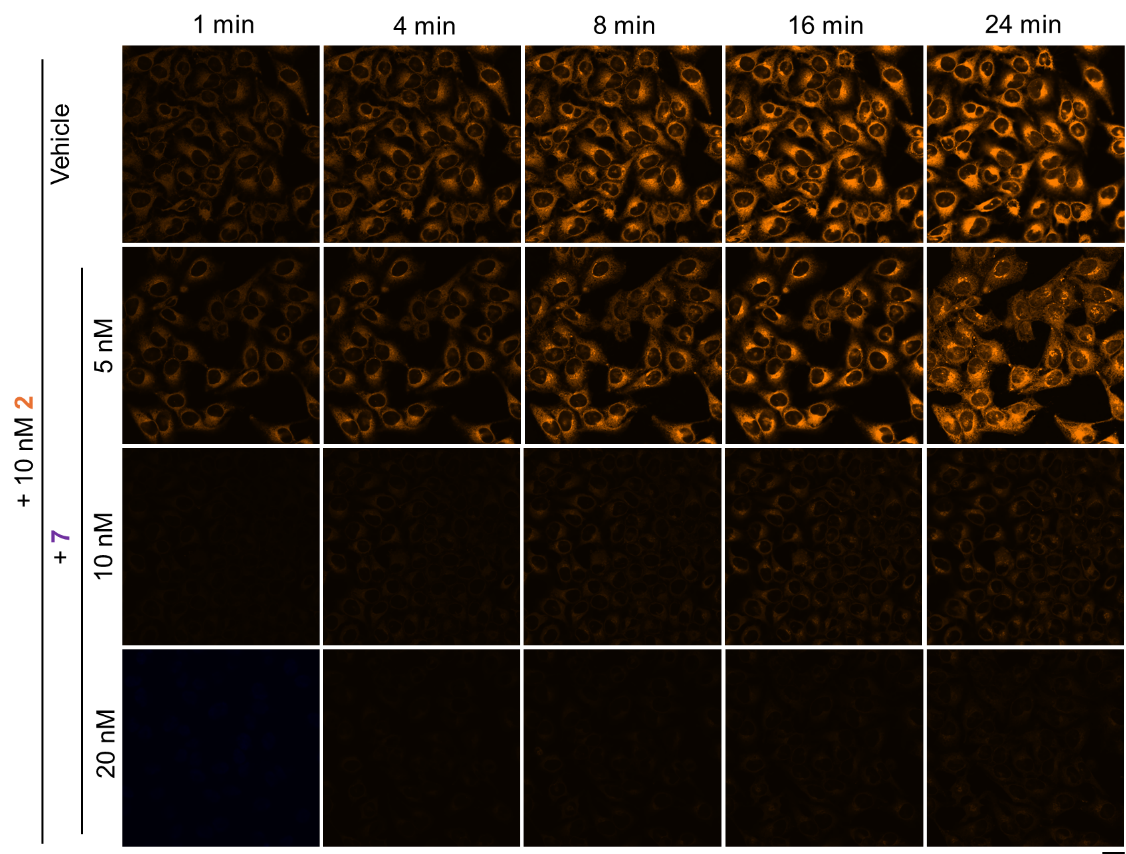

**Figure S13.** CLSM of HeLa cells were treated with **2** (10 nM) combined with **7** (5 nM, 10 nM, 20 nM) over 24 minutes. Scale bar = 20 μm.

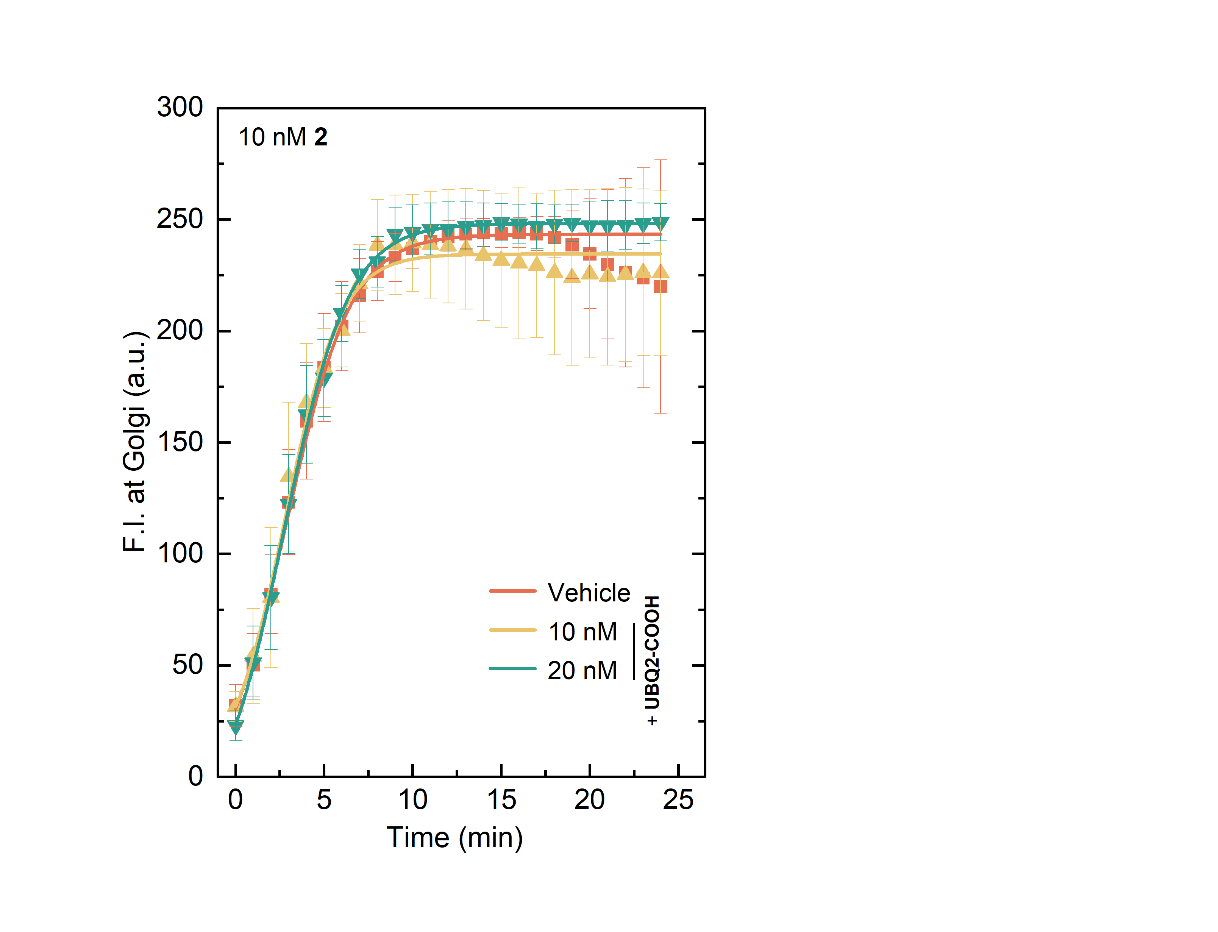

**Figure S14.** Golgi fluorescence intensity of HeLa cells treated with mixture of **2** (10 nM) and UBQ2-COOH (10, 20 nM) in 24 minutes.

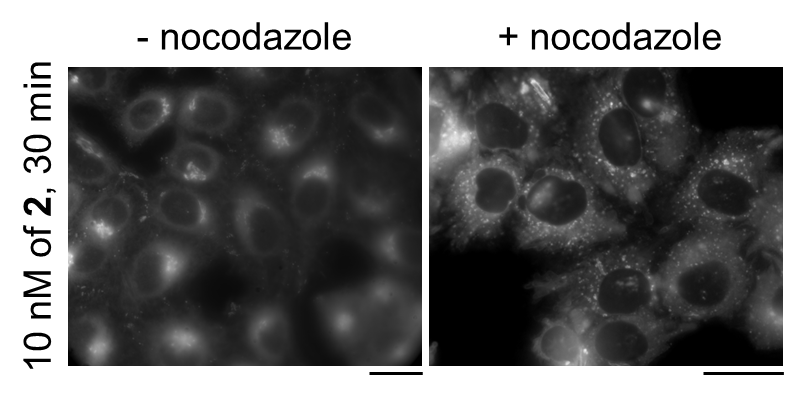

**Figure S15.** Widefield fluorescence microscopy of HeLa cells pretreated with vehicle or nocodazole (33 μM, 3 h) and then treated with **2** (10 nM) for 30 minutes. Scale bar = 20 μm.

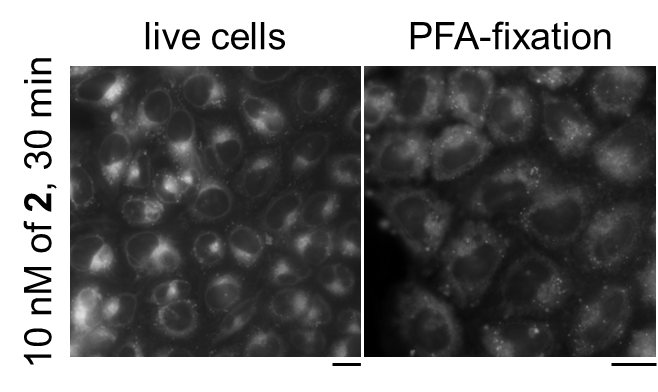

**Figure S16.** Widefield fluorescence microscopy of HeLa cells treated with **2** (10 nM) for 30 minutes, prior to fixation with 4% PFA. Scale bar = 20 μm.

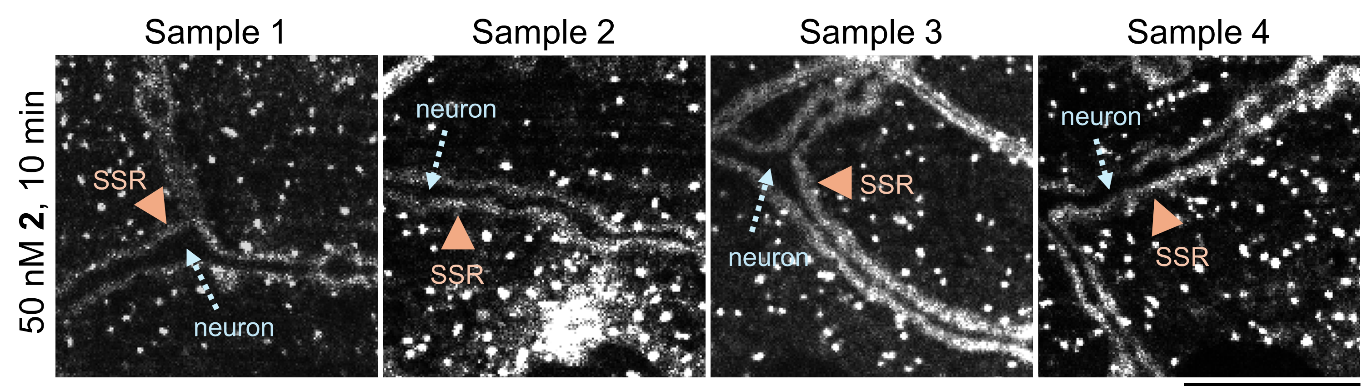

**Figure S17.** SDCM of Drosophila larval muscle treated with **2** (50 nM, 10 min). SSR is marked by orange arrows, inside of which are neurons. Fluorescent puncta are the Golgi. Scale bar = 20 μm.

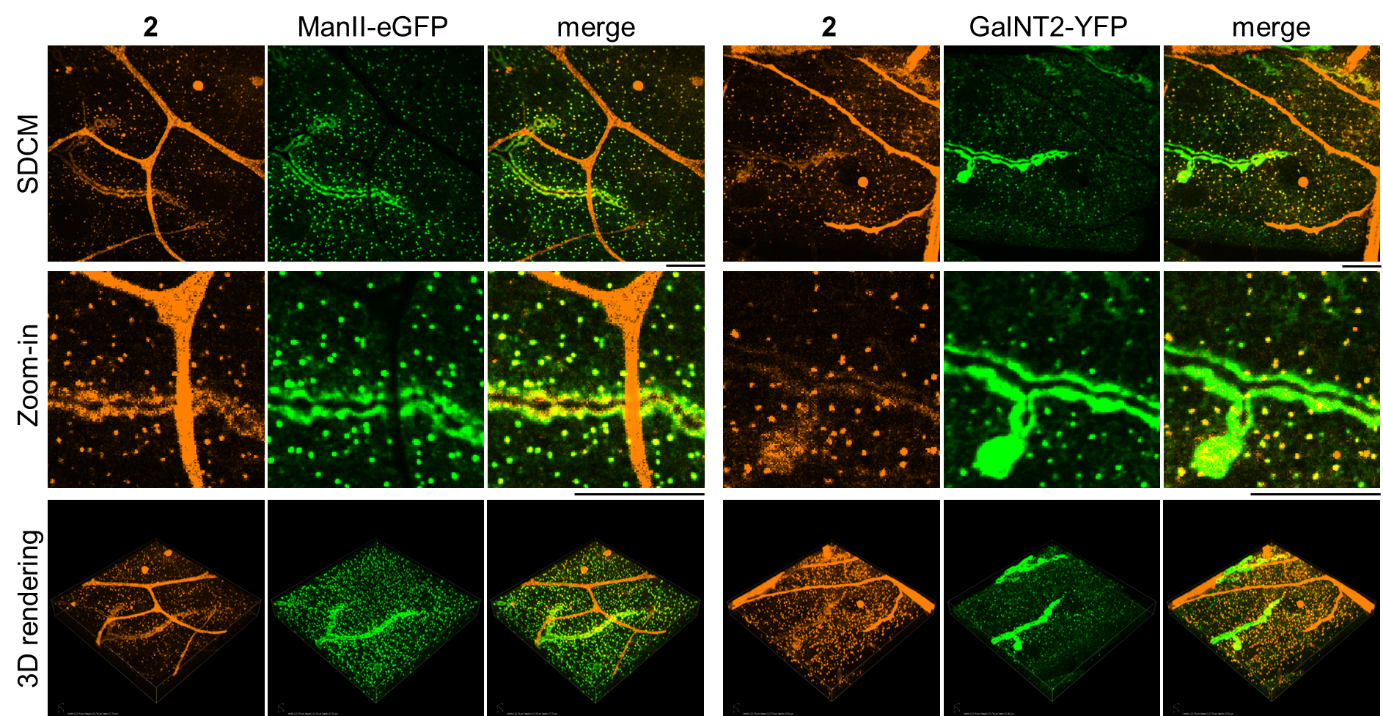

**Figure S18.** SDCM, zoom-in and 3D-rendering of *Drosophila* larval muscle treated with **2** (50 nM, 10 min) subsequent washout (30 min), with Golgi structures labeled using specific markers, ManII-eGFP (medial-Golgi) or GalNT2-YFP (trans-Golgi). Scale bar = 20 μm.

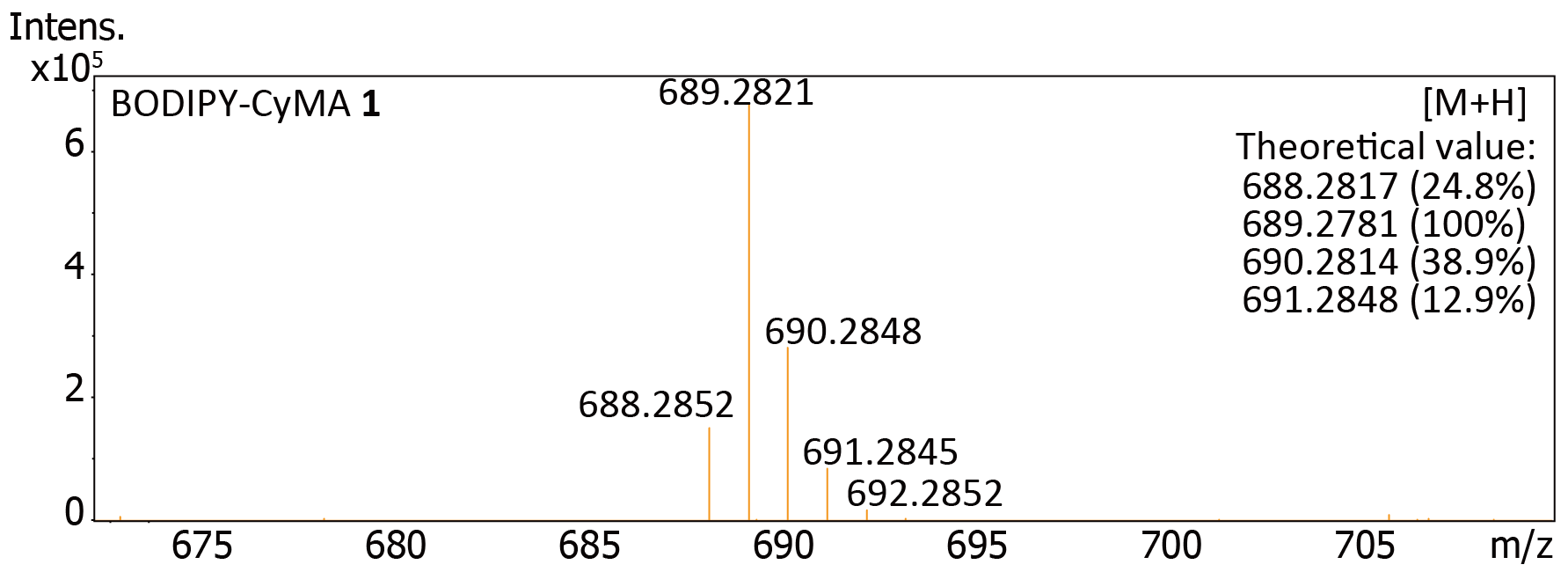

**Figure S19.** HRMS of BODIPY-CyMA **1**.

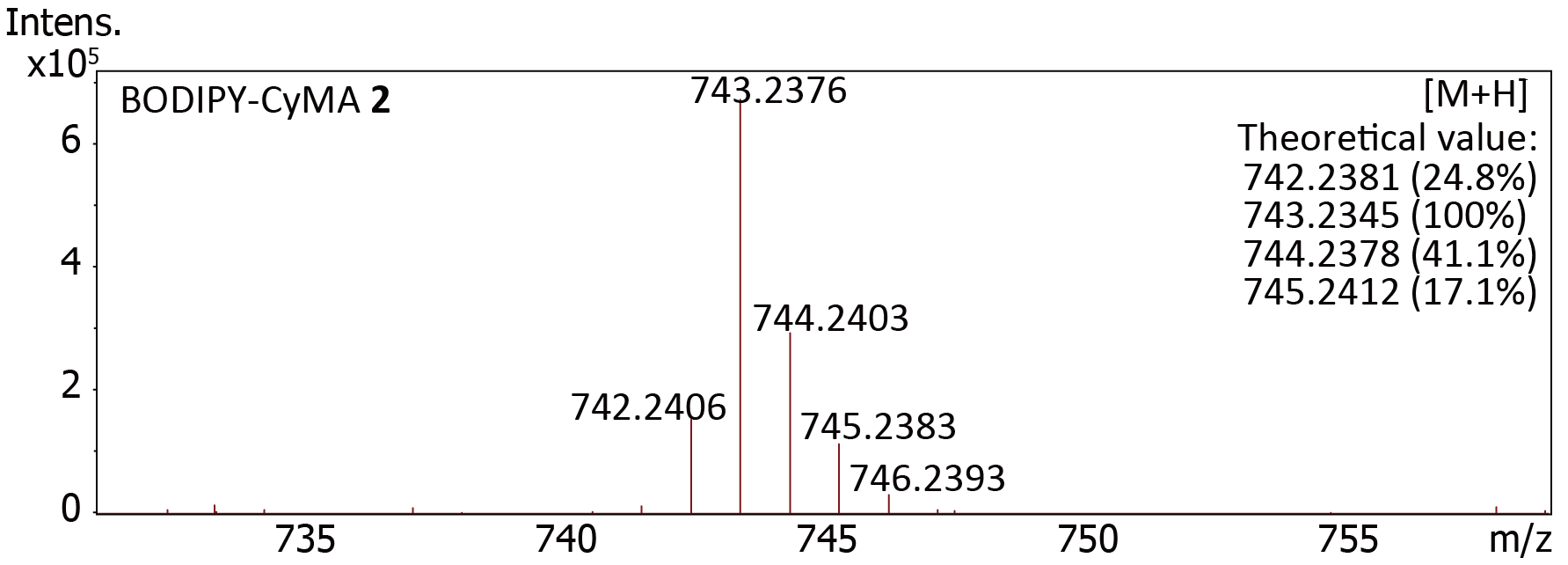

**Figure S20.** HRMS of BODIPY-CyMA **2**.

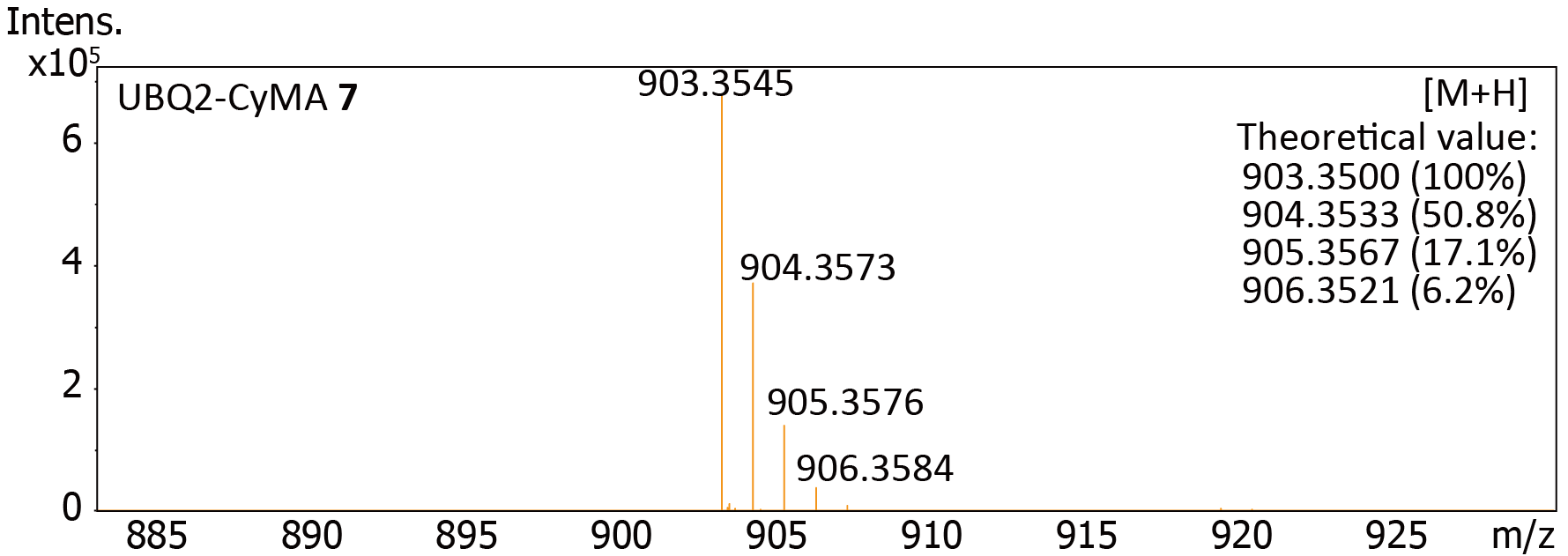

**Figure S21.** HRMS of UBQ2-CyMA **7**.

**Movie S1.** Time-series CLSM of HeLa cells treated with **1** or **2** (10 nM) over 30 minutes.

Writing:

The content and text were not generated by an LLM, but the grammar was proofread using LLM software.
